# Laser Flash Melting Cryo-EM Samples to Overcome Preferred Orientation

**DOI:** 10.1101/2024.11.21.624652

**Authors:** Monique S. Straub, Oliver F. Harder, Nathan J. Mowry, Sarah V. Barrass, Jakub Hruby, Marcel Drabbels, Ulrich J. Lorenz

## Abstract

Sample preparation remains a bottleneck for protein structure determination by cryo-electron microscopy. A frequently encountered issue is that proteins adsorb to the air-water interface of the sample in a limited number of orientations. This makes it challenging to obtain high-resolution reconstructions or may even cause projects to fail altogether. We have previously observed that laser flash melting and revitrification of cryo samples reduces preferred orientation for large, symmetric particles. Here, we demonstrate that our method can in fact be used to scramble the orientation of proteins of a range of sizes and symmetries. The effect can be enhanced for some proteins by increasing the heating rate during flash melting or by depositing amorphous ice onto the sample prior to revitrification. This also allows us to shed light onto the underlying mechanism. Our experiments establish a set of tools for overcoming preferred orientation that can be easily integrated into existing workflows.

---

Since the resolution revolution, cryo-electron microscopy (cryo-EM) has greatly expedited the process of protein structure determination ^1,2^. Advances in instrumentation and single-particle analysis now routinely enable high-resolution reconstructions for a wide range of proteins and protein complexes ^3–6^. However, sample preparation issues have remained a significant bottleneck ^7^. In particular, one frequently encountered challenge involves preferred particle orientation ^8–10^. When the sample solution is applied to the cryo-EM specimen support, the hydrophobic parts of the protein surface adsorb to the air-water interface, so that only a limited number of viewing directions are present after vitrification. Strong preferred orientation limits the resolution of a single-particle reconstruction along some viewing directions or may even make it difficult to obtain a reconstruction at all ^11^.

Although several strategies have been established to address issues with preferred orientation, no universal solution has been found. Tilting the sample provides a wider range of particle views, but usually also results in larger beam-induced motion, an increased ice thickness in the viewing direction, as well as a defocus gradient across the viewing area ^12,13^. Preferred orientation can be reduced by covering the air-water interface with detergents or small proteins in order to block adsorption of the particles of interest ^14,15^. Moreover, different specimen supports such as graphene, graphene oxide, or other functionalized surfaces can prevent some particles from diffusing to the interface ^16–20^. However, adopting such strategies often requires a time-consuming reoptimization of the sample preparation procedure. Adsorption to the air-water interface can also be reduced by limiting the time between sample application onto the specimen support and vitrification ^21–26^. While this time can be decreased to a few milliseconds, the timescale for the particles to diffuse to the interface is typically much shorter, on the order of a few microseconds. Significantly higher preparation speeds would therefore be required to prevent the majority of particles from reaching the interface. Finally, machine learning techniques have been introduced to guess missing views ^27–29^. However, such approaches cannot replace actual experimental observations.

We have previously observed that flash melting and revitrification of cryo-EM samples with microsecond laser pulses reduces preferred orientation for some large, highly-symmetric particles ^30–32^. Flash melting appears to exert small forces on the proteins, which changes their orientation and causes them to adopt an uneven spatial distribution after revitrification, with particles clustering together in some areas ^30,32^. However, it is not clear whether this may provide a practical tool for addressing preferred orientation or what the mechanism of the process is. Here, we show that this effect extends to smaller proteins as well as particles of lower symmetry. We also introduce two variants of the experiment that provide larger changes in preferred orientation for some proteins and that allow us to elucidate the underlying mechanism.

## Results and Discussion

### Flash melting with rectangular laser pulses

Our experimental approach is illustrated in Fig. 1a ^30,32–34^. A cryo-EM sample is flash melted with 20 µs or 30 µs rectangular laser pulses, *i*.*e*. the laser power is kept constant for the duration of the pulse (Fig. 1b). This causes some particles to detach from the air-water interface and change their orientation. Once the laser beam is switched off, the sample cools within just a few microseconds and revitrifies ^34,35^, trapping the proteins in a non-equilibrium angular distribution (Fig. 1a). *In situ* flash melting experiments are performed as previously described with a transmission electron microscope that we have modified for time-resolved experiments ^36,37^. The sample is locally flash melted by aiming the laser beam (532 nm wavelength, 28 µm diameter spot size in the sample plane) at the center of a grid square of the holey gold specimen support (1.2 µm holes, 1.3 µm apart on 300 mesh gold). Typically, an area of 9–16 grid holes is revitrified. The samples are subsequently transferred to a high-resolution electron microscope for imaging (Supplementary Information 1–4, 6).

**Figure 1.**
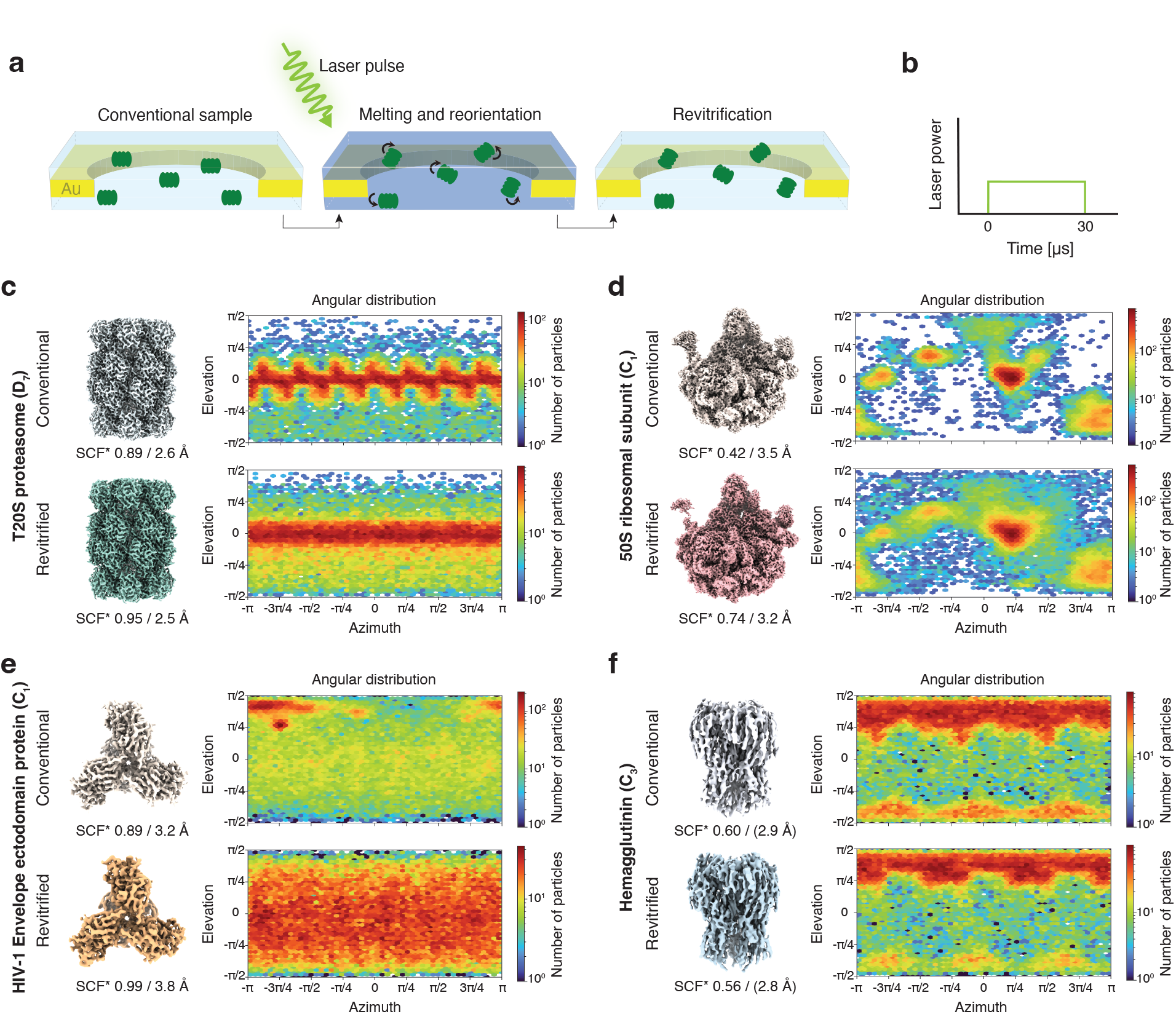
Revitrification with rectangular laser pulses. **a** Concept. A cryo-EM sample is flash melted with a 30 µs laser pulse, which scrambles the orientation of the particles. When the laser is switched off, the sample revitrifies, trapping the particles in a non-equilibrium angular distribution. **b** Schematic illustration of the rectangular laser pulse shape used. **c,d,e,f** Reconstructions, with the applied symmetry noted in parentheses, and angular distributions of conventional and revitrified samples (top and bottom, respectively) for the T20S proteasome (**c**), 50S ribosomal subunit (**d**), HIV-1 Envelope ectodomain protein (**e**) and Hemagglutinin (**f**). The sampling compensation factor (SCF*), which is a measure of the degree of preferred orientation, as well as the resolution of the reconstructions are indicated. Note that for Hemagglutinin, the resolution appears to be overestimated, as judged by the quality of the map.

Figure 1c demonstrates that flash melting reduces preferred orientation for a sample of the T20S proteasome. A reconstruction from a conventional sample is shown in the top, together with the angular distribution of the particles. It exhibits a series of maxima that arise from a strong preference of the T20S proteasome to adsorb to the air-water interface with its side, so that top views are only sparsely populated. Upon revitrification (bottom), the angular distribution broadens noticeably, with a larger fraction of the particles now populating the less favored views. At the same time, the sampling compensation factor (SCF*), a measure of the degree of preferred orientation, improves from 0.89 to 0.95. The SCF* takes values between 0 and 1, with 1 corresponding to a perfectly isotropic angular distribution, and SCF* values below 0.81 indicating strong preferred orientation ^5,38,39^.

An even more significant reduction in preferred orientation can be achieved for the 50S ribosomal subunit. Unlike the highly symmetric T20S proteasome (point group D_7_), the asymmetric 50S ribosomal subunit (C_1_) exhibits strong preferred orientation (Fig. 1d). In a conventional sample, the angular distribution features a small number of pronounced maxima, with many views absent altogether (white) and a low SCF* of 0.42. Upon revitrification, most of the missing views become populated, and the SCF* improves to 0.74. At the same time, the streaky artefacts that are visible in the reconstruction from the conventional sample and that are the result of preferred orientation disappear (Supplementary Information 2, Fig. S6) ^40^. Flash melting can also improve the angular distribution of much smaller proteins, such as the HIV-1 Envelope ectodomain protein (210 kDa), for which the SCF* increases from 0.89 to 0.99 (Fig. 1e). Note that among these examples, a noticeable gain in resolution is only observed for the 50S ribosomal subunit, the only protein for which preferred orientation is so pronounced that many views are not populated at all. For the HIV-1 Envelope ectodomain protein, the resolution slightly decreases. This seems to be largely due to astigmatism present in the micrographs recorded for the revitrified sample.

Laser flash melting appears to be less efficient at reducing preferred orientation for proteins that are more strongly bound to the air-water interface. Hemagglutinin (170 kDa) exhibits a preference for top and bottom views (Fig. 1f). While revitrification somewhat depopulates the bottom views, the overall angular distribution remains similar, with the SCF* slightly decreasing from 0.60 to 0.56. This suggests either that flash melting detaches only few proteins from the interface or that detached particles return to their preferred orientation rapidly once they have diffused back to the surface. Either explanation points to a strong interaction of Hemagglutinin with the air-water interface. Note that for the proteins studied here, diffusion to the interface is expected to occur in just a few microseconds, which is shorter than the duration of the experiment (30 µs).

### Flash melting with shaped laser pulses

Experiments with shaped microsecond laser pulses provide clues about the mechanism by which the angular distribution of the particles is reshuffled. We have previously shown that cryo-EM samples partially crystallize during flash melting with rectangular laser pulses ^34,41^. This has led us to speculate whether the transient formation of crystallites, which preferentially form at the sample surface ^42^, could exert small forces on the particles that cause them to detach and reorient ^30^. We test this hypothesis by melting samples with shaped laser pulses that feature an intense initial spike of 1 µs duration and ten times the laser power (Fig. 2a,b) ^43^. This increases the heating rate during flash melting to about 2·10^8^ K/s, twice the critical heating rate of 10^8^ K/s, so that the sample does not crystallize ^43^. If crystallization was the dominant mechanism for reshuffling the particle orientations, we would expect the angular distribution to barely change under these conditions. Contrary to our expectation, flash melting with such a shaped laser pulse decreases preferred orientation even more drastically for the 50S ribosomal subunit (Fig. 2c, Supplementary Information 2). The SCF* increases from 0.18 to 0.90 in revitrified areas of the sample, and the map resolution improves significantly, from 4.1 Å to 2.9 Å. Note that the angular distribution of the sample is somewhat different from that in Fig. 1d, even though both samples were prepared with the same procedure. This suggests that the interaction of the particles with the interface is very sensitive to the preparation conditions and can be easily altered.

**Figure 2.**
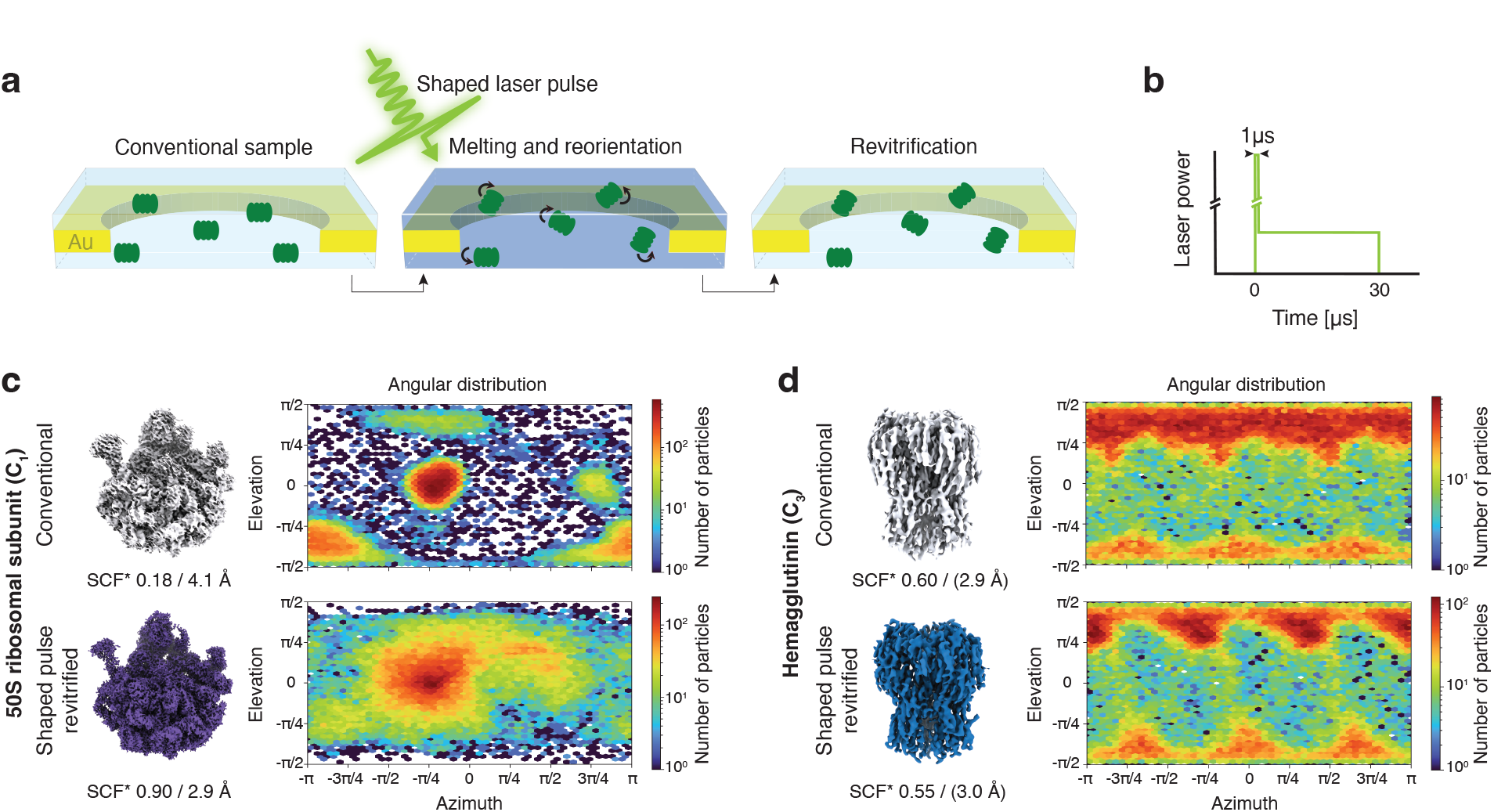
Revitrification with shaped laser pulses featuring an intense leading edge. **a** Concept. A cryo-EM sample is flash melted with a shaped 30 µs laser pulse that features an intense leading edge. When the laser is switched off, the sample revitrifies, with the angular distribution of the particles reshuffled. **b** Schematic illustration of the laser pulse shape, which features an intense spike at its onset (1 µs duration with the laser power increased ten-fold). **c,d** Reconstructions and angular distributions of conventional and revitrified samples (top and bottom, respectively) for the 50S ribosomal subunit (**c**) and Hemagglutinin (**d**). The sampling compensation factor (SCF*) and the resolution of the reconstructions are shown below the maps. Note that for Hemagglutinin, the resolution appears to be overestimated, judging by the quality of the map.

Clearly, transient crystallization of the sample cannot be the dominant mechanism for reorienting the particles. Instead, the effect appears to become more pronounced as the heating rate is increased, which suggests several possible mechanisms. It is well known that impulsive laser heating of a thin, suspended membrane, such as the holey gold film of the specimen support, induces drumming motions that typically persist on microsecond timescales ^44,45^. With the larger initial heating power of the shaped laser pulses, such oscillations likely reach a larger amplitude and exert greater forces on the thin liquid film in which the particles are suspended. This may allow them to detach from the air-water interface more readily and reorient. Note that the impinging laser beam also deforms the thin liquid film directly through the radiation forces that act upon it ^46^. Finally, it is also conceivable that some particles detach from the interface due to the evaporation of the liquid sample in the vacuum of the electron microscope ^41^. During the initial spike of the shaped laser pulse, the sample temperature briefly overshoots by about 25 K, as simulations reveal (Supplementary Information 5). Since the evaporation rate increases exponentially with temperature ^47^, this could cause more particles to detach. Note however, that we also observe changes in preferred orientation when we revitrify samples at atmospheric pressure, where evaporation is significantly reduced ^48,49^. It is therefore unlikely that evaporation is the dominant effect that causes particles to detach from the air-water interface. For Hemagglutinin, even shaped laser pulses are not able to reduce the amount of preferred orientation, although they induce some changes in the angular distribution (Fig. 2d, Supplementary Information 4).

### Flash melting after deposition of a layer of amorphous ice

Our experiments suggest that the changes in the angular distribution induced by flash melting are the result of two opposing processes. First, particles must be successfully detached from the air-water interface in order to reorient. But given enough time, they may then diffuse back to the interface and return to their preferred orientation. A simple experiment allows us to detach all particles from the interface before we laser melt the sample and thus disentangle both processes (Fig. 3a). We deposit a 20 nm thick layer of amorphous ice onto both sides of the sample by dosing water vapor into the volume surrounding the specimen in our modified transmission electron microscope ^50^. Once we flash melt the sample with a rectangular laser pulse (Fig. 3b), the particles all start out at a distance from the interface and should have lost all memory of where they used to be located. The angular distribution obtained after revitrification should then exclusively reflect the result of the free diffusion of the particles.

**Figure 3.**
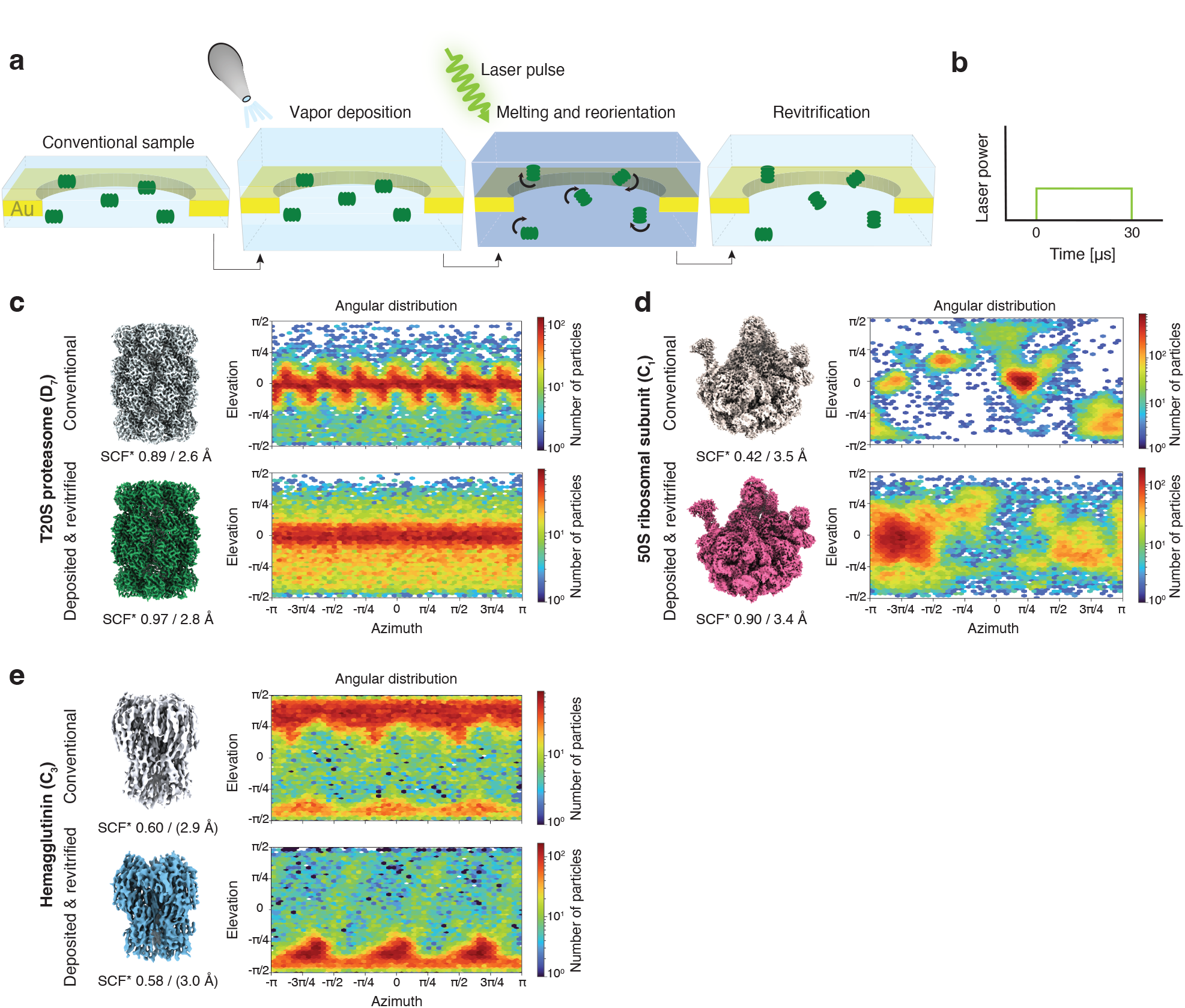
Revitrification with rectangular laser pulses after deposition of amorphous ice. **a** Concept. A 20 nm layer of amorphous ice is deposited *in situ* onto the cryo-EM sample before it is flash melted with a 30 µs laser pulse. When the laser is switched off, the sample revitrifies with the particle orientations redistributed. **b** Schematic illustration of the rectangular laser pulse shape used. **c,d,e** Reconstructions and angular distributions of conventional and revitrified samples (top and bottom, respectively) for the T20S proteasome (**c**), the 50S ribosomal subunit (**d**), and Hemagglutinin (**e**). The sampling compensation factor (SCF*) and the resolution of the reconstructions are indicated. Note that for Hemagglutinin, the resolution appears to be overestimated, judging by the quality of the map.

When we revitrify a T20S proteasome sample after depositing a layer of amorphous ice, we obtain the angular distribution in Fig. 3c (bottom, Supplementary Information 1). With the estimated rotational timescale of the particles (about 5 µs) shorter than the timescale of the experiment, one might naïvely expect that this should yield an isotropic distribution. Instead, the distribution closely resembles that obtained without prior deposition of amorphous ice (Fig. 1c) and notably features a preference for side views. This clearly indicates that the particles have diffused back to the interface, which occurs on an estimated timescale of only about 4 µs, and have partially settled into their preferred orientations. The striking similarity of the angular distribution to that in Fig. 1c suggests that both are the result of the free diffusion and partial readsorption of the particles. It therefore appears likely that even without prior deposition of amorphous ice, the laser pulse efficiently detaches the T20S proteasome particles from the interface.

Flash melting of a cryo-EM sample of the 50S ribosomal subunit after deposition of amorphous ice similarly results in a broad angular distribution with a small degree of preferred orientation, indicating that some particles have readsorbed to the interface (Fig. 3d, Supplementary Information 2). The SCF* has increased from 0.42 to 0.90. Curiously however, the distribution has significantly changed and now features maxima at orientations that were previously only sparsely populated. This suggests that the interaction of the particles with the interface has been altered in our experiments. As discussed above, the preferential orientation of the 50S ribosomal subunit varies even in conventional samples under otherwise identical preparation conditions (Figs. 1d and 2c), suggesting that the properties of the interface are easily modified. It is conceivable that sample transfer and handling have introduced traces of surface-active compounds and thus changed the chemical composition of the interface. It is also possible that the interaction of the particles with the surface has been altered due to the dilution of the buffer that occurs in our experiment. At the laser powers used, less material is evaporated than has been deposited, so that the sample thickness slightly increases overall. As shown in Fig. 3e, we obtain a similar result for Hemagglutinin (Supplementary Information 4). After amorphous ice deposition and revitrification, the particles show a significant amount of preferred orientation, indicating that many have readsorbed to the interface. However, the prominent top views have been largely depopulated, while the bottom views now dominate the angular distribution.

## Conclusion

Our experiments yield a simple picture of the competing processes that occur during laser revitrification. Flash melting detaches some particles from the interface (Fig. 4a,b), a process that becomes more efficient as the heating rate is increased. As these particles rotate freely, their orientations are scrambled, which leads to an improvement of the angular distribution. In a second, competing process, particles diffuse back to the interface (Fig. 4c) before the sample is revitrified (Fig. 4d), which allows some proteins to efficiently readsorb in their preferred orientation. For such proteins, it may be possible to improve the angular distribution by reducing the duration of the laser pulse to below the timescale of diffusion (a few microseconds), so that the particles do not have enough time to reach the interface. In some experiments, we observe that proteins readsorb in a different preferred orientation, such as when we deposit a layer of amorphous ice prior to revitrification. Apparently, our experiment has altered the interfacial properties, so that adsorption in new orientations has become favorable. This also suggests that it should be possible to purposely modify the interface prior to flash melting, for example through vapor deposition of a hydrophilic compound onto the cryo-EM sample. Upon laser melting, all particles should then desorb and randomize their orientation. Such experiments are currently underway in our lab.

**Figure 4.**
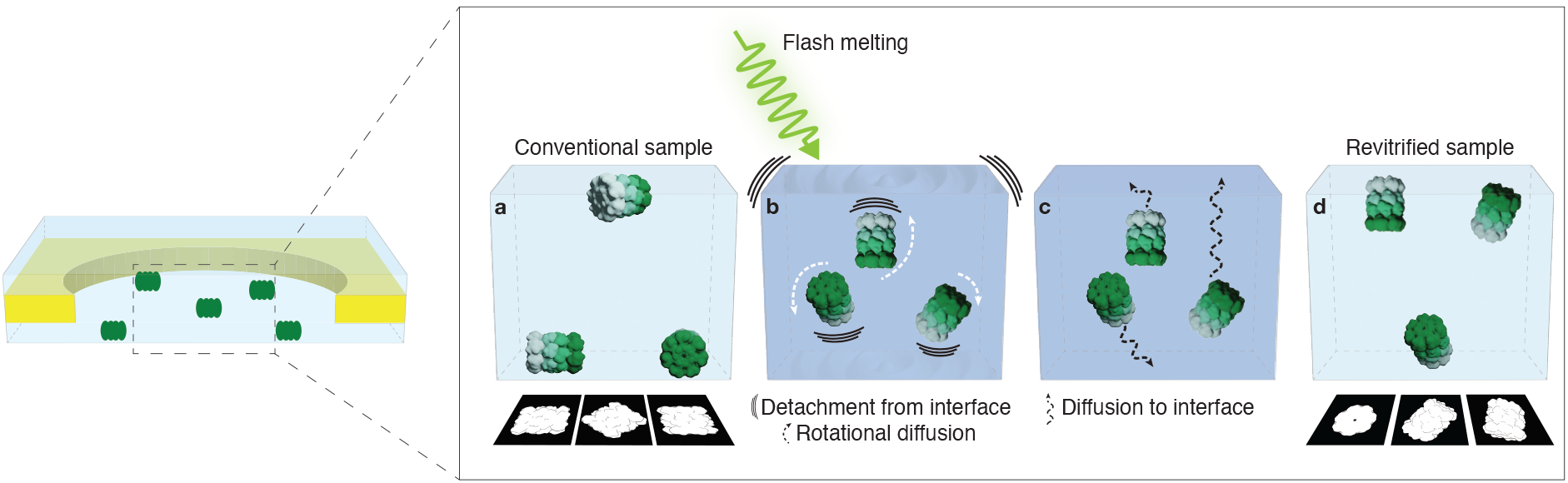
Competing processes determine the degree to which laser flash melting is able to scramble the particle orientations. **a** In a conventional cryo-EM sample, particles adsorb to the air-water interface in preferred orientations. **b** Flash melting detaches the proteins from the interface, so that they are able to rotate freely. **c** While the sample is liquid, many particles diffuse back to the interface, where some adsorb in their preferred orientation. **d** The sample is revitrified with the ensemble arrested in a non-equilibrium angular distribution.

Our experiments also provide a practical toolbox for addressing preferred orientation, with some proteins showing dramatic improvements in their angular distribution. For example, revitrifying a cryo-EM sample of the 50S ribosomal subunit with shaped laser pulses improves the resolution by 1.2 Å under otherwise identical conditions (Fig. 2c). To put this in perspective, it would be necessary to acquire 18 times as many micrographs from a conventional sample to achieve the same improvement in resolution. Reshuffling the particle orientations through laser flash melting can therefore provide significant cost and time savings. We note that our method can be easily combined with other established approaches for addressing preferred orientation. Importantly, as a simple physical approach, flash melting does not require any time-consuming changes to the sample preparation procedure, but can be easily integrated into existing workflows. As we have previously shown, cryo-EM samples can be revitrified in an optical microscope equipped for correlative light and electron microscopy experiments, an approach that may be most easily accessible to other labs ^48^. It is also conceivable that laser melting experiments could be directly integrated into high-resolution electron microscopes at cryo-EM facilities. If a sample is found to exhibit preferred orientation during data acquisition, it could then be revitrified on the fly with little additional time added to the workflow.

## Methods

### Cryo-EM sample preparation

Purified T20S Proteasome (3.7 mg/ml in 50 mM Tris pH 8.0, 200 mM sodium chloride) and purified 50S Ribosome ^51^ (40 OD_260_/ml in 20 mM HEPES pH 7.5, 100 mM sodium chloride, 2 mM magnesium chloride) were gifted by A. Guskov, University of Gröningnen, NL, and B. Beckert, DCI Lausanne, CH, respectively. The purified Hemagglutinin sample (1mg/ml in PBS pH 7.5) was purchased from MyBioSource (MBS434205). The cryo-EM samples of purified proteins were prepared by applying 3.5 µl of the sample solution to glow-discharged (0.25 mbar, 15 mA, 60 s) gold grids (UltrAuFoil, R1.2/1.3, 300 mesh, Quantifoil Micro Tools GmbH). Surplus sample was removed by blotting for 2 to 4 s using a blot force of 10 in a Vitrobot Mark IV (Thermo Fisher Scientific, held at 20°C, 100% relative humidity). The HIV-1 Envelope ectodomain protein (HIV-1 CH505TF Envelope SOSIP ectodomain) ^52^ was generously provided by R. Henderson, Duke Center for HIV Structural Biology, US. The HIV-1 Envelope ectodomain protein (5 mg/ml in 15 mM HEPES pH 7.4, 150 mM NaCl, 2.5% glycerol) was applied to glow discharged gold grids (Au-flat, 0.6 µm/1 µm, 300 mesh, Electron Microscopy Sciences). The sample was blotted with the minimum blot time that can be set (0 s). The conventional sample was blotted using a blot force of 10, whereas the revitrified sample was blotted using a blot force of 5 and 10. All samples were plunge-frozen in liquid ethane and stored in liquid nitrogen until experiments were carried out.

### Flash melting with rectangular laser pulses

The revitrification experiments with rectangular laser pulses (Fig. 1) were performed *in situ* using a modified JEOL 2200FS transmission electron microscope equipped with a TemCam-XF416 (TVIPS) and an in-column Omega energy filter ^37^. The samples were loaded into the microscope using a single-tilt side-entry cryo-transfer holder (Elsa, Gatan) held at approximately 100 K. The sample was melted and revitrified in the center of multiple grid squares (typically 80 – 100) through irradiation with microsecond laser pulses as previously described ^30,32,34^. A 30 µs laser pulse was used for the 50S and HIV-1 samples, and a 20 μs laser pulse for the T20S sample. The Hemagglutinin sample was revitrified with two 30 µs pulses with a 250 ms delay between the pulses. The laser pulses were obtained by chopping the continuous output of a 532 nm laser (Ventus532, Laser Quantum) with an acousto-optic modulator (AA Opto-Electronic). The laser beam was focused to a spot size of 28 μm FWHM (full width at half maximum) in the sample plane, as determined by a knife-edge scan, and aligned to the center of the grid squares. The laser power was adjusted on the fly to keep the size of the revitrified region, typically 15 – 22 µm in diameter, approximately constant ^34^. After revitrification, the grids were unloaded from the side entry holder and stored in liquid nitrogen until data collection.

### Flash melting with shaped laser pulses

The revitrification experiments with shaped laser pulses (Fig. 2) were performed on a modified JEOL 2010F transmission electron microscope, equipped with a TemCam-XF416 (TVIPS) ^50,53,54^. The plunge-frozen cryo-EM samples were inserted into the microscope using a single-tilt side-entry cryo-transfer holder (Elsa, Gatan) held at a temperature of approximately 100 K. Shaped laser pulses with an intense leading edge (1 µs initial spike at ten times the laser power) were generated as described previously ^43^ by modulating the output of a continuous green laser (532 nm, Verdi Coherent) with an acousto-optic modulator (AA Opto-Electronic) that was controlled by an arbitrary function generator (Tektronix AFG1062). The laser beam was focused to a spot of 38 µm FWHM in the sample plane, as determined by imaging the beam with CCD camera placed in a plane that is conjugate to the sample plane. The laser power required for laser melting was initially determined by using ordinary rectangular pulses. The initial spike was then added to perform the revitrification experiment.

### Flash melting with rectangular laser pulses after deposition of amorphous ice

The experiments in Fig. 3, in which the sample was revitrified after deposition of amorphous ice, were performed on a modified JEOL 2010F transmission electron microscope, equipped with a TemCam-XF416 (TVIPS) ^41,50,53,54^. Amorphous ice was deposited by leaking water vapor into the volume surrounding the sample through a gas dosing valve, as described previously ^50^. In order to obtain a well-defined thickness, the deposition rate was first determined with the following procedure. A multilayer graphene sample on a gold specimen grid (UltrAu Foil, 600 mesh, R2/1) was cooled to about 100 K in an Elsa cryo-transfer holder (Gatan). While amorphous ice was being deposited on the graphene sheet, diffraction patterns of the growing ice layer were recorded. During the deposition, the intensity of the molecular diffraction pattern of amorphous ice initially increased linearly but leveled off and went through a maximum once the ice film was so thick that multiple and inelastic scattering became important. From the characteristic curve shape of the diffraction intensity as a function of deposition time, the deposition rate was then determined (typically 0.3 nm/s) ^41,50^, and the deposition time was adjusted accordingly (typically 74 s to deposit about 10 nm on each side of the sample, 20 nm total). In order to perform the deposition and revitrification experiment, the cryo-EM sample was loaded into the microscope with a second Elsa cryo-transfer holder (Gatan, approximately at 100 K). To begin the deposition, the shutter of the specimen holder was opened for the predetermined amount of time, before the shutter was closed to stop any further deposition of amorphous ice. The gas dosing valve was then closed, and the cryo shield surrounding the sample was cooled to liquid nitrogen temperature. Once the pressure in the microscope column had dropped to below 5·10^−5^ Pa, the shutter of the cryo-transfer holder was opened, and the revitrification experiment was performed using rectangular laser pulses, as described above.

## Acknowledgements

The authors would like to thank B. Beckert (Dubochet Center for Imaging Lausanne, CH), A. Guskov (University of Gröningen, NL), and R. Henderson (Duke University, US) for providing the protein samples used in this study. A. Myasnikov, B. Beckert, S. Nazarov, I. Mohammed and E. Uchikawa, the members of the Dubochet Center for Imaging in Lausanne, and P. Szwedziak at the Center for Microscopy and Image Analysis in Zurich are acknowledged for assistance with cryo-EM data collection. The authors would like to acknowledge current and former members of the lab for their help at various stages of the project.

## Author Contributions

These authors contributed equally: Monique S. Straub, Oliver F. Harder, Nathan J. Mowry

UJL was responsible for conceptualizing this work. The methodology was performed by MSS, OFH, NJM, SVB, and JH. MSS, OFH, NJM, SVB, and JH performed cryo-EM sample preparation. MSS, SVB and JH acquired the cryo-EM data. MSS and SVB performed cryo-EM data processing. NJM performed the simulations. MSS, MD, and UJL performed the data analysis. Acquiring funding, project administration, and supervision was performed by UJL. The writing of the original draft and data visualization was performed by MSS and UJL. The reviewing and editing of the manuscript were performed by MSS and UJL with the input of all co-authors. The work was supported by Swiss National Science Foundation Grant TMCG-2_213773 and by the Duke Center for HIV Structural Biology, NIH grant U54AI170752.

## Materials & Correspondence

### Supplementary Information is available for this paper

The data that support the findings of this study are available from the corresponding author upon request. The cryo-EM maps have been deposited in the Electron Microscopy Data Bank (EMDB) and the Electron Microscopy Public Image Archive (EMPIAR) under accession codes EMD-51744 and EMPIAR-12389 (T20S conventional), EMD-51745 and EMPIAR-12388 (T20S revitrified), EMD-51746 and EMPIAR-12390 (T20S revitrified after deposition), EMD-51747 and EMPIAR-12397 (50S conventional), EMD-51748 and EMPIAR-12398 (50S revitrified), EMD-51749 and EMPIAR-12399 (50S revitrified after deposition), EMD-51750 and EMPIAR-12435 (50S shaped pulse conventional), EMD-51751 and EMPIAR-12436 (50S shaped pulse revitrified), EMD-51752 and EMPIAR-12437 (HIV conventional), EMD-51753 and EMPIAR-12438 (HIV revitrified), EMD-51754 and EMPIAR-12439 (HA conventional), EMD-51755 and EMPIAR-12440 (HA revitrified), EMD-51756 and EMPIAR-12441 (HA revitrified after deposition), and EMD-51757 and EMPIAR-12442 (HA shaped pulse revitrified).

## References

1. Hand, E. Cheap shots. Science 367, 354–358 (2020).

2. Chua, E. Y. D. et al. Better, Faster, Cheaper: Recent Advances in Cryo–Electron Microscopy. Annu. Rev. Biochem. 91, 1–32 (2022).

3. Nakane, T. et al. Single-particle cryo-EM at atomic resolution. Nature 587, 152–156 (2020).

4. Yip, K. M., Fischer, N., Paknia, E., Chari, A. & Stark, H. Atomic-resolution protein structure determination by cryo-EM. Nature 587, 157–161 (2020).

5. Punjani, A., Rubinstein, J. L., Fleet, D. J. & Brubaker, M. A. cryoSPARC: algorithms for rapid unsupervised cryo-EM structure determination. Nat. Methods 14, 290–296 (2017).

6. Scheres, S. H. W. RELION: Implementation of a Bayesian approach to cryo-EM structure determination. J. Struct. Biol. 180, 519–530 (2012).

7. Weissenberger, G., Henderikx, R. J. M. & Peters, P. J. Understanding the invisible hands of sample preparation for cryo-EM. Nat. Methods 18, 463–471 (2021).

8. Taylor, K. A. & Glaeser, R. M. Retrospective on the early development of cryoelectron microscopy of macromolecules and a prospective on opportunities for the future. J. Struct. Biol. 163, 214–223 (2008).

9. Noble, A. J. et al. Routine single particle CryoEM sample and grid characterization by tomography. eLife 7, e34257 (2018).

10. Glaeser, R. M. & Han, B.-G. Opinion: hazards faced by macromolecules when confined to thin aqueous films. Biophys. Rep. 3, 1–7 (2017).

11. Drulyte, I. et al. Approaches to altering particle distributions in cryo-electron microscopy sample preparation. Acta Crystallogr. Sect. Struct. Biol. 74, 560–571 (2018).

12. Tan, Y. Z. et al. Addressing preferred specimen orientation in single-particle cryo-EM through tilting. Nat. Methods 14, 793–796 (2017).

13. Aiyer, S. et al. Overcoming resolution attenuation during tilted cryo-EM data collection. Nat. Commun. 15, 389 (2024).

14. Abe, K. M., Li, G., He, Q., Grant, T. & Lim, C. J. Small LEA proteins mitigate air-water interface damage to fragile cryo-EM samples during plunge freezing. Nat. Commun. 15, 7705 (2024).

15. Chen, J., Noble, A. J., Kang, J. Y. & Darst, S. A. Eliminating effects of particle adsorption to the air/water interface in single-particle cryo-electron microscopy: Bacterial RNA polymerase and CHAPSO. J. Struct. Biol. X 1, 100005 (2019).

16. Glaeser, R. M. et al. Factors that Influence the Formation and Stability of Thin, Cryo-EM Specimens. Biophys. J. 110, 749–755 (2016).

17. Han, Y. et al. High-yield monolayer graphene grids for near-atomic resolution cryoelectron microscopy. Proc. Natl. Acad. Sci. 117, 1009–1014 (2020).

18. D’Imprima, E. et al. Protein denaturation at the air-water interface and how to prevent it. eLife 8, e42747 (2019).

19. Cookis, T. et al. Streptavidin-Affinity Grid Fabrication for Cryo-Electron Microscopy Sample Preparation. https://app.jove.com/t/66197/streptavidin-affinity-grid-fabrication-for-cryo-electron-microscopy-sample-preparation.

20. Zheng, L. et al. Self-assembled superstructure alleviates air-water interface effect in cryo-EM. Nat. Commun. 15, 7300 (2024).

21. Klebl, D. P. et al. Need for Speed: Examining Protein Behavior during CryoEM Grid Preparation at Different Timescales. Structure 28, 1238-1248.e4 (2020).

22. Dandey, V. P. et al. Time-resolved cryo-EM using Spotiton. Nat. Methods 17, 897–900 (2020).

23. Rima, L. et al. cryoWriter: a blotting free cryo-EM preparation system with a climate jet and cover-slip injector. Faraday Discuss. 240, 55–66 (2022).

24. Yang, Z. et al. Electrospray-assisted cryo-EM sample preparation to mitigate interfacial effects. Nat. Methods 21, 1023–1032 (2024).

25. H.J.V. Tyrrell, K.R. Harris. Diffusion in Liquids-A Theoretical and Experimental Study. (Butterworth-Heinemann, 1984).

26. Mäeots, M.-E. & Enchev, R. I. Structural dynamics: review of time-resolved cryo-EM. Acta Crystallogr. Sect. Struct. Biol. 78, 927–935 (2022).

27. Zhang, Zheng et al. Addressing preferred orientation in single-particle cryo-EM through AI-generated auxiliary particles. BioRxiv Prepr. Serv. Biol. (2023).

28. Liu, Y.-T., Hu, J. & Zhou, Z. H. Resolving the Preferred Orientation Problem in CryoEM Reconstruction with Self-Supervised Deep Learning. Microsc. Microanal. 29, 1918–1919 (2023).

29. Liu, Y.-T., Fan, H., Hu, J. J. & Zhou, Z. H. Overcoming the preferred-orientation problem in cryo-EM with self-supervised deep learning. Nat. Methods 1–11 (2024)

30. Bongiovanni, G., Harder, O. F., Voss, J. M., Drabbels, M. & Lorenz, U. J. Near-atomic resolution reconstructions from in situ revitrified cryo samples. Acta Crystallogr. Sect. Struct. Biol. 79, 473– 478 (2023).

31. Lorenz, U. J. Microsecond time-resolved cryo-electron microscopy. Curr. Opin. Struct. Biol. 87, 102840 (2024).

32. Harder, O. F., Barrass, S. V., Drabbels, M. & Lorenz, U. J. Fast viral dynamics revealed by microsecond time-resolved cryo-EM. Nat. Commun. 14, 5649 (2023).

33. U.J. Lorenz, G. Bongiovanni, O.F. Harder, J.M. Voss, M. Drabbels, M. Straub, N.J. Mowry. Methods to overcome preferred orientation in cryo-samples for single particle analysis - provisional patent application.

34. Voss, J. M., Harder, O. F., Olshin, P. K., Drabbels, M. & Lorenz, U. J. Microsecond melting and revitrification of cryo samples. Struct. Dyn. 8, 054302 (2021).

35. Mowry, N. J., Kruger, C. R., Drabbels, M. & Lorenz, U. J. Direct Measurement of the Critical Cooling Rate for the Vitrification of Water. Preprint at 10.48550/arXiv.2407.01087 (2024).

36. Russo, C. J. & Passmore, L. A. Ultrastable gold substrates for electron cryomicroscopy. Science 346, 1377–1380 (2014).

37. Olshin, P. K., Drabbels, M. & Lorenz, U. J. Characterization of a time-resolved electron microscope with a Schottky field emission gun. Struct. Dyn. 7, 054304 (2020).

38. Baldwin, P. R. & Lyumkis, D. Non-uniformity of projection distributions attenuates resolution in Cryo-EM. Prog. Biophys. Mol. Biol. 150, 160–183 (2020).

39. Baldwin, P. R. & Lyumkis, D. Tools for visualizing and analyzing Fourier space sampling in Cryo-EM. Prog. Biophys. Mol. Biol. 160, 53–65 (2021).

40. Lander, G. C. Single particle cryo-EM map and model validation: It’s not crystal clear. Curr. Opin. Struct. Biol. 89, 102918 (2024).

41. Mowry, N. J., Krüger, C. R., Bongiovanni, G., Drabbels, M. & Lorenz, U. J. Flash melting amorphous ice. J. Chem. Phys. 160, 184502 (2024).

42. Backus, E. H. G., Grecea, M. L., Kleyn, A. W. & Bonn, M. Surface Crystallization of Amorphous Solid Water. Phys. Rev. Lett. 92, 236101 (2004).

43. Krüger, C. R., Mowry, N. J., Drabbels, M. & Lorenz, U. J. Shaped Laser Pulses for Microsecond Time-Resolved Cryo-EM: Outrunning Crystallization during Flash Melting. J. Phys. Chem. Lett. 15, 4244–4248 (2024).

44. Kwon, O.-H., Barwick, B., Park, H. S., Baskin, J. S. & Zewail, A. H. Nanoscale Mechanical Drumming Visualized by 4D Electron Microscopy. Nano Lett. 8, 3557–3562 (2008).

45. Lorenz, U. J. & Zewail, A. H. Biomechanics of DNA structures visualized by 4D electron microscopy. Proc. Natl. Acad. Sci. 110, 2822–2827 (2013).

46. Astrath, N. G. C., Malacarne, L. C., Baesso, M. L., Lukasievicz, G. V. B. & Bialkowski, S. E. Unravelling the effects of radiation forces in water. Nat. Commun. 5, 4363 (2014).

47. Robert J. List. Smithsonian Meteorological Tables. 114, 539 (1968).

48. Bongiovanni, G., Harder, O. F., Drabbels, M. & Lorenz, U. J. Microsecond melting and revitrification of cryo samples with a correlative light-electron microscopy approach. Front. Mol. Biosci. 9, (2022).

49. Marek, R. & Straub, J. Analysis of the evaporation coefficient and the condensation coefficient of water. Int. J. Heat Mass Transf. 44, 39–53 (2001).

50. Krüger, C. R., Mowry, N. J., Bongiovanni, G., Drabbels, M. & Lorenz, U. J. Electron diffraction of deeply supercooled water in no man’s land. Nat. Commun. 14, 2812 (2023).

51. Paternoga, H. et al. Structural conservation of antibiotic interaction with ribosomes. Nat. Struct. Mol. Biol. 30, 1380–1392 (2023).

52. LaBranche, C. C. et al. Neutralization-guided design of HIV-1 envelope trimers with high affinity for the unmutated common ancestor of CH235 lineage CD4bs broadly neutralizing antibodies. PLOS Pathog. 15, e1008026 (2019).

53. Olshin, P. K., Bongiovanni, G., Drabbels, M. & Lorenz, U. J. Atomic-Resolution Imaging of Fast Nanoscale Dynamics with Bright Microsecond Electron Pulses. Nano Lett. 21, 612–618 (2021).

54. Bongiovanni, G., Olshin, P. K., Drabbels, M. & Lorenz, U. J. Intense microsecond electron pulses from a Schottky emitter. Appl. Phys. Lett. 116, 234103 (2020).

