## Supplementary Information for "Laser Flash Melting Cryo-EM Samples to Overcome Preferred Orientation"

**This PDF file includes:**

1 | Data collection and analysis – T20S proteasome

2 | Data collection and analysis – 50S ribosomal subunit

3 | Data collection and analysis – HIV-1 Envelope ectodomain protein

4 | Data collection and analysis – Hemagglutinin

5 | Simulation of the temperature evolution of the sample

6 | Cryo-EM data collection, refinement and validation statistics

7 | References

### 1 | Data collection and analysis – T20S proteasome

The T20S datasets were recorded on a Titan Krios G3i, equipped with a BioQuantum energy filter and a K3 camera, at the Center for Microscopy and Image Analysis in Zurich, Switzerland. The movies were recorded as tiff images in super-resolution mode and binned on the fly to a pixel size of 0.651 Å. The energy filter slit was set to 20 eV. The data were collected using a 100 µm objective aperture at a nominal magnification of 130’000 x during a 1.3 s exposure with a defocus range of -0.6 µm to -2.0 µm and a total dose of 67 $e^{-}/Å^{2}$.

The datasets were processed in cryoSPARC v4.4 and v4.5 [1]. For the conventional sample, the revitrified sample, and the sample revitrified after deposition of amorphous ice, 3’479, 6’105, and 3’641 micrographs were collected, respectively. The micrographs were subjected to patch motion-correction and patch CTF estimation prior to manual curation based on CTF resolution estimation, ice thickness and total full-frame motion. Particles were initially picked with a blob picker with a diameter of 100–200 Å from 1’519, 2’287, and 2’255 high quality micrographs, respectively, and were then extracted with a box size of 512 px and down sampled to 128 px. After two rounds of 2D classifications, the selected particles were used for *ab initio* reconstruction with two classes. The particles assigned to the better resolved class were refined against the obtained volume in a homogeneous refinement with $C_1$ symmetry. The resulting volume was used to generate templates for template-based picking with a particle diameter of 150 Å. The particles were again extracted with a box size of 512 px, Fourier-cropped to a box size of 256 px, and subjected to two rounds of 2D classification. The selected particles were used for *ab initio* reconstruction with one class only, followed by homogeneous refinement using $D_7$ symmetry. Subsequently, 50’000 particles were randomly selected, re-extracted at full box size and again directed into a homogeneous refinement with $D_7$ symmetry. To ensure the comparability of the datasets obtained from the different experimental conditions, a low pass filtered volume (filtered to 30 Å) obtained from the conventional dataset was used as a reference input for homogeneous refinements for all the three datasets. Finally, an orientation diagnostics job in cryoSPARC was run to assess the angular distribution of the particles.

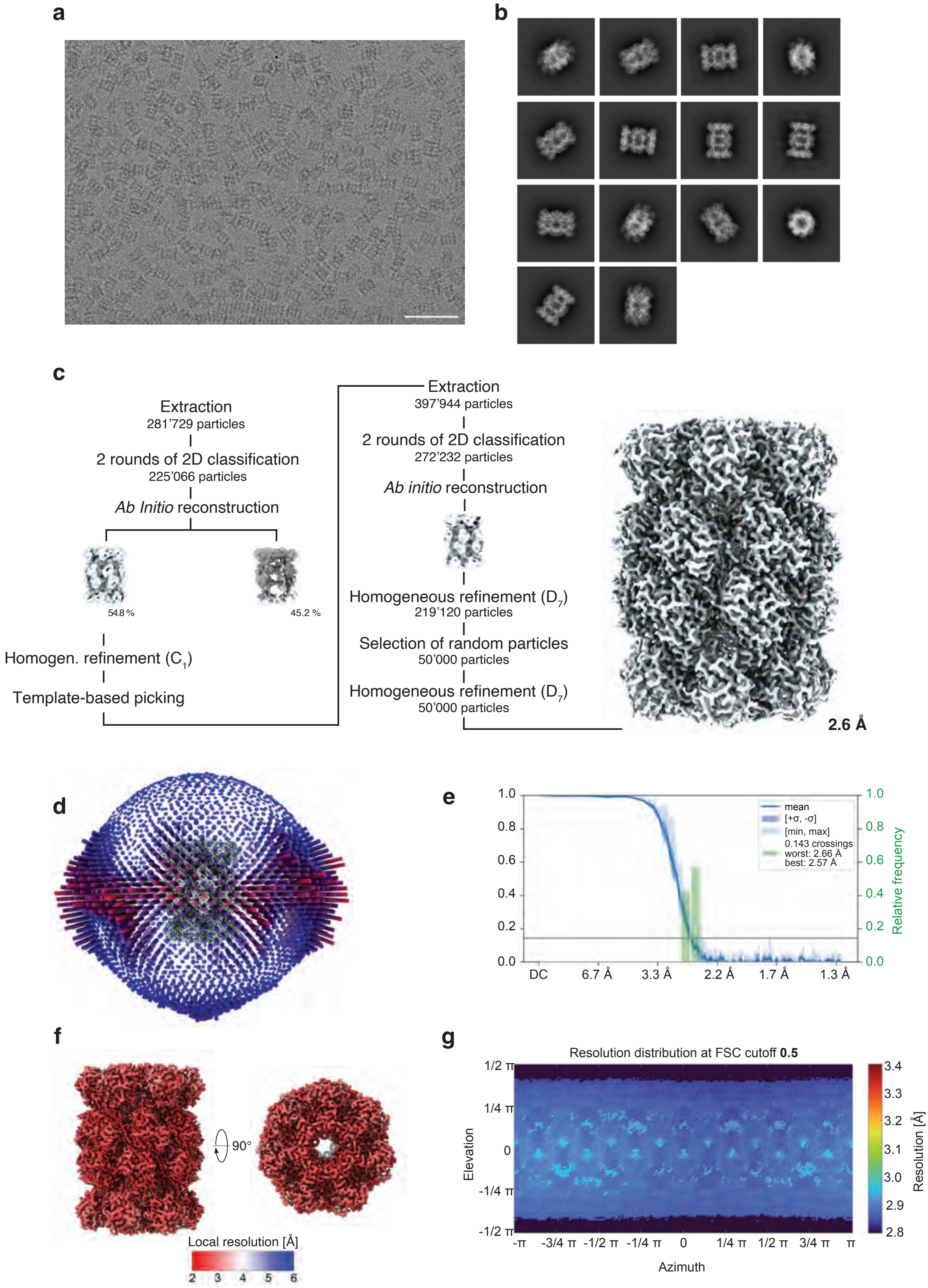
a
b
c
Extraction
281’729 particles
2 rounds of 2D classification
225’066 particles
Ab Initio reconstruction
54.8 %
45.2 %
Homogen. refinement ($C_1$)
Template-based picking
Extraction
397’944 particles
2 rounds of 2D classification
272’232 particles
Ab initio reconstruction
Homogeneous refinement ($D_7$)
219’120 particles
Selection of random particles
50’000 particles
Homogeneous refinement ($D_7$)
50’000 particles
2.6 Å
d
e
mean
[+σ, -σ]
[min, max]
0.143 crossings
worst: 2.66 Å
best: 2.57 Å
Relative frequency
DC
6.7 Å
3.3 Å
2.2 Å
1.7 Å
1.3 Å
f
90°
Local resolution [Å]
2 3 4 5 6
g
Resolution distribution at FSC cutoff **0.5**
Elevation
Azimuth
Resolution [Å]
-π -3/4 π -1/2 π -1/4 π 0 1/4 π 1/2 π 3/4 π π

**Supplementary Figure 1 | Cryo-EM data processing workflow for a conventional sample of the T20S proteasome. a** Representative micrograph, revealing that side views of the proteasome are predominantly obtained. Scale bar, 500 Å. **b** 2D class averages, underlining the abundance of side views. **c** Data processing workflow in cryoSPARC. The symmetry applied in each step is indicated in parentheses. A resolution of 2.6 Å was obtained in the final reconstruction. The map is shown at a 5 $\sigma$. **d** Three-dimensional representation of angular distribution of the particles, with the color and height of the cylinders representing the frequency of observation (red: more abundant, blue: less abundant). **e** Plot of the 3DFSC with the 0.143 cutoff indicated (grey line). The mean FSC value is shown as a solid blue line, with dark blue shading corresponding to one standard deviation, and the light blue shading indicating the minimum and maximum values. The green bars represent a histogram of the resolution values obtained from the directional FSC. **f** Final map with the local resolution estimation indicated in color. **g** The resolution distribution plot shown at FSC cutoff of 0.5, as obtained from the 3DFSC job in cryoSPARC.

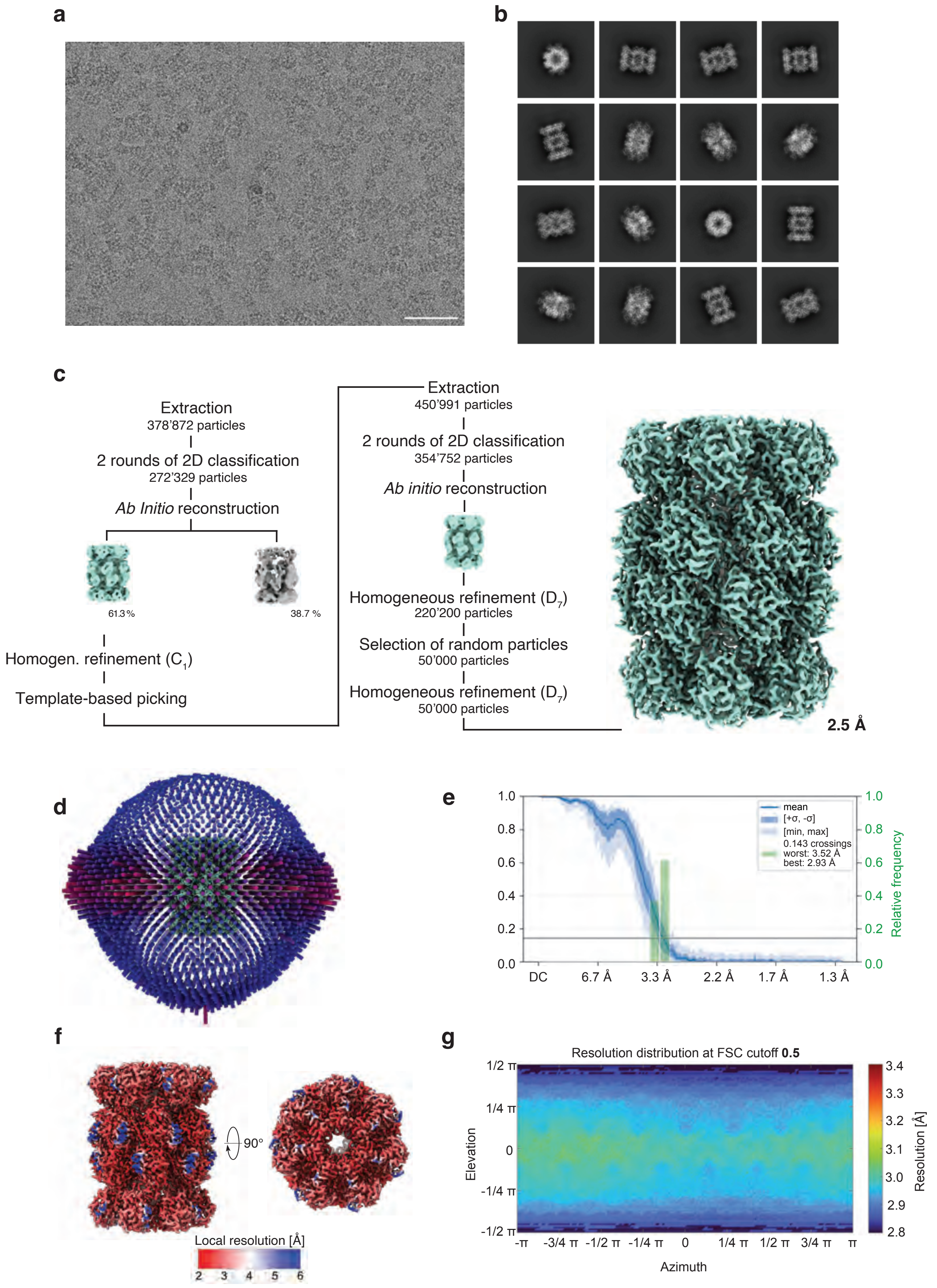
a
b
c
Extraction
378’872 particles
2 rounds of 2D classification
272’329 particles
Ab Initio reconstruction
61.3 %
38.7 %
Homogen. refinement (C1)
Template-based picking
Extraction
450’991 particles
2 rounds of 2D classification
354’752 particles
Ab initio reconstruction
Homogeneous refinement (D7)
220’200 particles
Selection of random particles
50’000 particles
Homogeneous refinement (D7)
50’000 particles
2.5 Å
d
e
mean
[+σ, -σ]
[min, max]
0.143 crossings
worst: 3.52 Å
best: 2.93 Å
Relative frequency
DC
6.7 Å
3.3 Å
2.2 Å
1.7 Å
1.3 Å
f
90°
Local resolution [Å]
2 3 4 5 6
g
Resolution distribution at FSC cutoff 0.5
Elevation
Azimuth
Resolution [Å]
1/2 π
1/4 π
0
-1/4 π
-1/2 π
-π
-3/4 π
-1/2 π
-1/4 π
1/4 π
1/2 π
3/4 π
π

**Supplementary Figure 2 | Cryo-EM data processing workflow for a revitrified sample of the T20S proteasome.** **a** Representative micrograph. Scale bar 500 Å. **b** 2D class averages revealing more tilted views of the T20S proteasome. **c** Data processing workflow in cryoSPARC. The symmetry applied is indicated in parentheses. A resolution of 2.5 Å was obtained in the final reconstruction. The map is shown 5 $\sigma$. **d** Three-dimensional representation of angular distribution of the particles, with the color and height of the cylinders representing the frequency of observation (red: more abundant, blue: less abundant). **e** Plot of the 3DFSC with the 0.143 cutoff indicated (grey line). The mean FSC value is shown as a solid blue line, with dark blue shading corresponding to one standard deviation, and the light blue shading indicating the minimum and maximum values. The green bars represent a histogram of the resolution values obtained from the directional FSC. **f** Final map with the local resolution estimation indicated in color. **g** The resolution distribution plot shown at FSC cutoff of 0.5, as obtained from the 3DFSC job in cryoSPARC. The directional resolutions appear slightly worse for the revitrified, compared to the conventional sample. However, the differences are minimal (within 0.1 Å) and are therefore considered as noise.

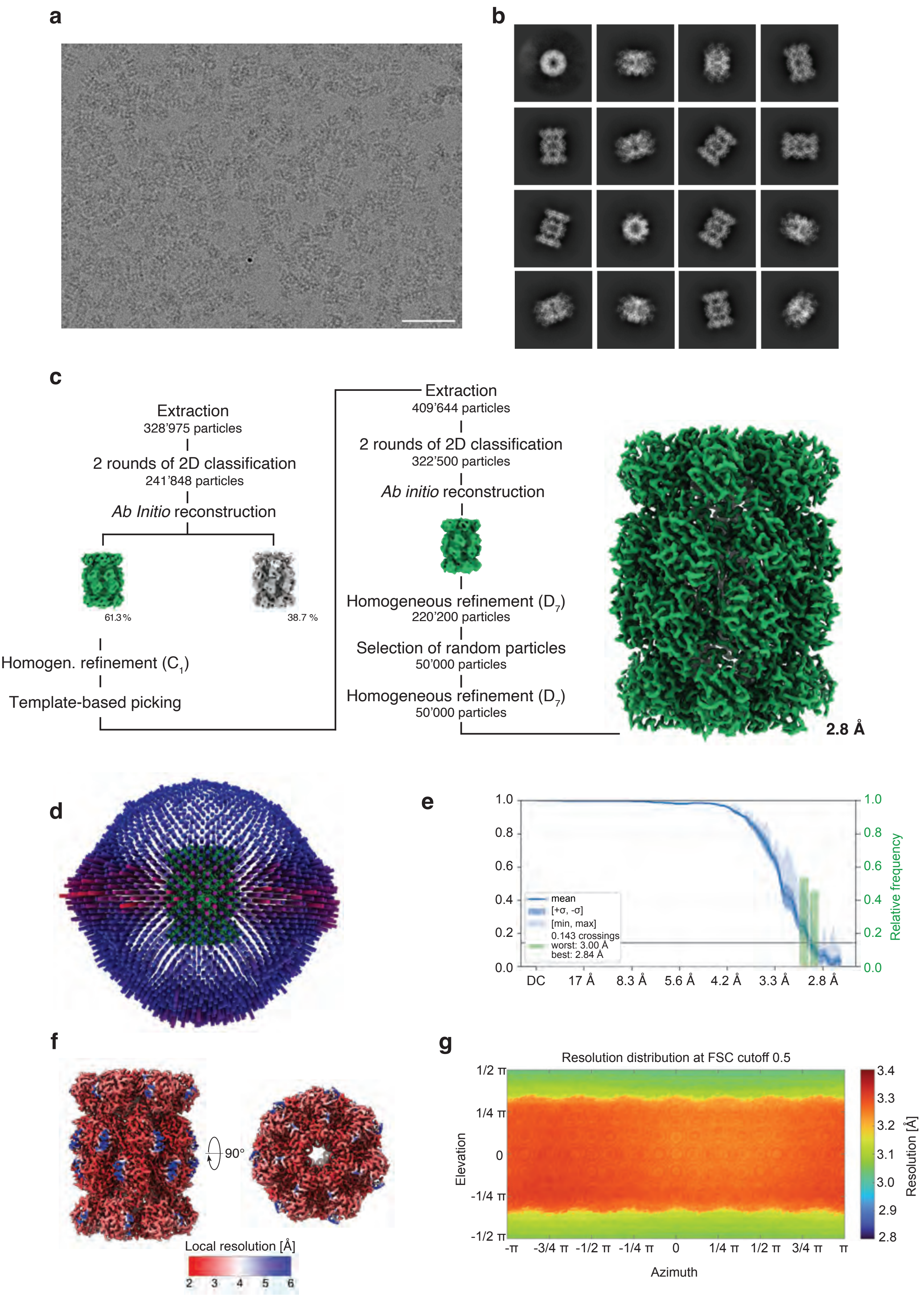
a
b
c
Extraction
328'975 particles
2 rounds of 2D classification
241'848 particles
*Ab Initio* reconstruction
61.3 %
38.7 %
Homogen. refinement ($C_1$)
Template-based picking
Extraction
409'644 particles
2 rounds of 2D classification
322'500 particles
*Ab initio* reconstruction
Homogeneous refinement ($D_7$)
220'200 particles
Selection of random particles
50'000 particles
Homogeneous refinement ($D_7$)
50'000 particles
2.8 Å
d
e
mean
[+σ, -σ]
[min, max]
0.143 crossings
worst: 3.00 Å
best: 2.84 Å
Relative frequency
DC
17 Å
8.3 Å
5.6 Å
4.2 Å
3.3 Å
2.8 Å
f
90°
Local resolution [Å]
2 3 4 5 6
g
Resolution distribution at FSC cutoff 0.5
Elevation
Azimuth
Resolution [Å]
1/2 π
1/4 π
0
-1/4 π
-1/2 π
-π
-3/4 π
-1/2 π
-1/4 π
1/4 π
1/2 π
3/4 π
π

**Supplementary Figure 3 | Cryo-EM data processing workflow for a deposited and revitrified sample of the T20S proteasome. a** Representative micrograph. Scale bar 500 Å. **b** 2D class averages, revealing more tilted views. **c** Data processing workflow in cryoSPARC. The symmetry applied in each step is indicated in parentheses. A resolution of 2.8 Å was obtained in the final reconstruction. The map is contoured 5 $\sigma$. **d** Three-dimensional representation of angular distribution of the particles, with the color and the height of the cylinders representing the frequency of observation (red: more abundant, blue: less abundant). **d** Plot of the 3DFSC with the 0.143 cutoff indicated (grey line). The mean FSC value is shown as a solid blue line, with dark blue shading corresponding to one standard deviation, and the light blue shading indicating the minimum and maximum values. The green bars represent a histogram of the resolution values obtained from the directional FSC. **f** Final map with the local resolution estimation indicated in color. **g** The resolution distribution plot shown at FSC cutoff of 0.5, as obtained from the 3DFSC job in cryoSPARC. Due to the increase in ice thickness after deposition, the directional resolutions, as well as the overall resolution of the reconstruction, decreased minimally.

### 2 | Data collection and analysis – 50S ribosomal subunit

The conventional, revitrified, and deposited and revitrified 50S datasets were collected on a Titan Krios G4 equipped with a Falcon 4 camera at the Dubochet Center for Imaging facility in Lausanne, Switzerland. The movies were collected in eer format, with a 100 µm objective aperture inserted, at a nominal magnification of 120’000 x, corresponding to a pixel size of 0.658 Å per pixel. Micrographs were collected with a defocus range of -0.4 µm to -1 µm using a total dose of 40 $e^-/Å^2$.

The sample revitrified with shaped laser pulses and the corresponding conventional control were collected on a Titan Krios G3i, fitted with a BioQuantum energy filter and a K3 camera, at the Center for Microscopy and Image Analysis in Zurich, Switzerland. The movies were recorded during a 1.2 s exposure at 130’000 x magnification, corresponding to a pixel size of 0.3255 Å in super-resolution mode. The data was binned on the fly to a pixel size of 0.651 Å per pixel. The energy filter slit was set to 20 eV and the objective aperture to 100 µm. The defocus used ranged from -0.4 µm to -1.6 µm, whereas the electron dose amounted 63 $e^-/Å^2$.

For the 50S datasets, 11’121, 6’174, 9’532, 7’416 and 7’725 micrographs were collected for conventional, revitrified, deposited and revitrified, conventional (control for shaped pulses) and shaped pulse revitrified, respectively. The micrographs were patch motion-corrected (40 fractions, no up-sampling), and patch CTF-estimated before being subjected to a manual curation, where low quality micrographs, judged by CTF estimation, relative ice thickness and total full-frame motion, were discarded. The selected 6’236, 5’870, 6’641, 3’485, and 5’227 high quality micrographs were subjected to a blob picker with a particle diameter of 200-300 Å. The particles were extracted with a box size of 784 px and cropped to 392 px. After two rounds of 2D classification, selected particles were directed into an *ab initio* reconstruction with two to three classes. The particles from the best class were refined against the volumes from the best and worst class in one round of heterogeneous refinement. The selected particles from the heterogeneous refinement were directed into a homogeneous refinement with $C_1$ symmetry. Subsequently, 50’000 particles were randomly selected, re-extracted at full box size, and refined in a homogeneous refinement using $C_1$ symmetry and a low pass filtered volume from the conventional dataset as a reference. Ultimately, an orientation diagnostics job in cryoSPARC was run to assess the angular distribution of the particles.

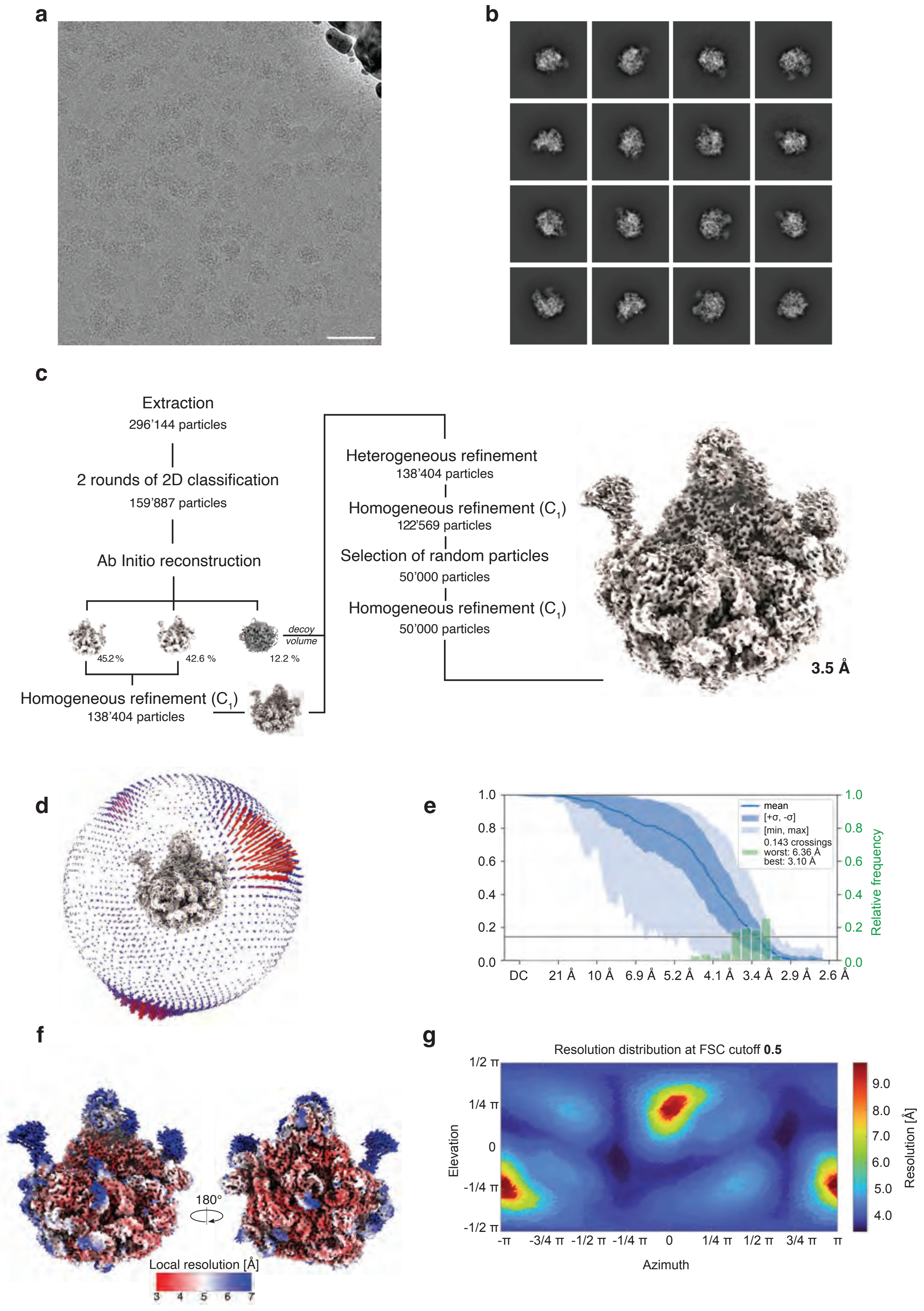
a
b
c
Extraction
296’144 particles
2 rounds of 2D classification
159’887 particles
Ab Initio reconstruction
45.2 %
42.6 %
12.2 %
decoy volume
Homogeneous refinement (C1)
138’404 particles
Heterogeneous refinement
138’404 particles
Homogeneous refinement (C1)
122’569 particles
Selection of random particles
50’000 particles
Homogeneous refinement (C1)
50’000 particles
3.5 Å
d
e
mean
[+σ, -σ]
[min, max]
0.143 crossings
worst: 6.36 Å
best: 3.10 Å
Relative frequency
DC 21 Å 10 Å 6.9 Å 5.2 Å 4.1 Å 3.4 Å 2.9 Å 2.6 Å
f
180°
Local resolution [Å]
3 4 5 6 7
g
Resolution distribution at FSC cutoff 0.5
Elevation
Azimuth
Resolution [Å]
-π -3/4 π -1/2 π -1/4 π 0 1/4 π 1/2 π 3/4 π π

**Supplementary Figure 4 | Cryo-EM data processing workflow for a conventional sample of the 50S ribosome.** **a** Representative micrograph. Scale bar 100 Å. **b** 2D class averages **c** Data processing workflow in cryoSPARC. The symmetry applied in each step is indicated in parentheses. A resolution of 3.5 Å was obtained in the final reconstruction. The map is shown at 2.8 σ. **d** Three-dimensional representation of angular distribution of the particles, with the color and the height of the cylinders representing the frequency of observation (red: more abundant, blue: less abundant). **d** Plot of the 3DFSC with the 0.143 cutoff indicated (grey line). The mean FSC value is shown as a solid blue line, with dark blue shading corresponding to one standard deviation, and the light blue shading indicating the minimum and maximum values. The green bars represent a histogram of the resolution values obtained from the directional FSC. **f** Final map with the local resolution estimation indicated in color. **g** The resolution distribution plot shown at FSC cutoff of 0.5, as obtained from the 3DFSC job in cryoSPARC. The plot shows regions with lower resolutions, corresponding to missing views in the angular distribution (panel d).

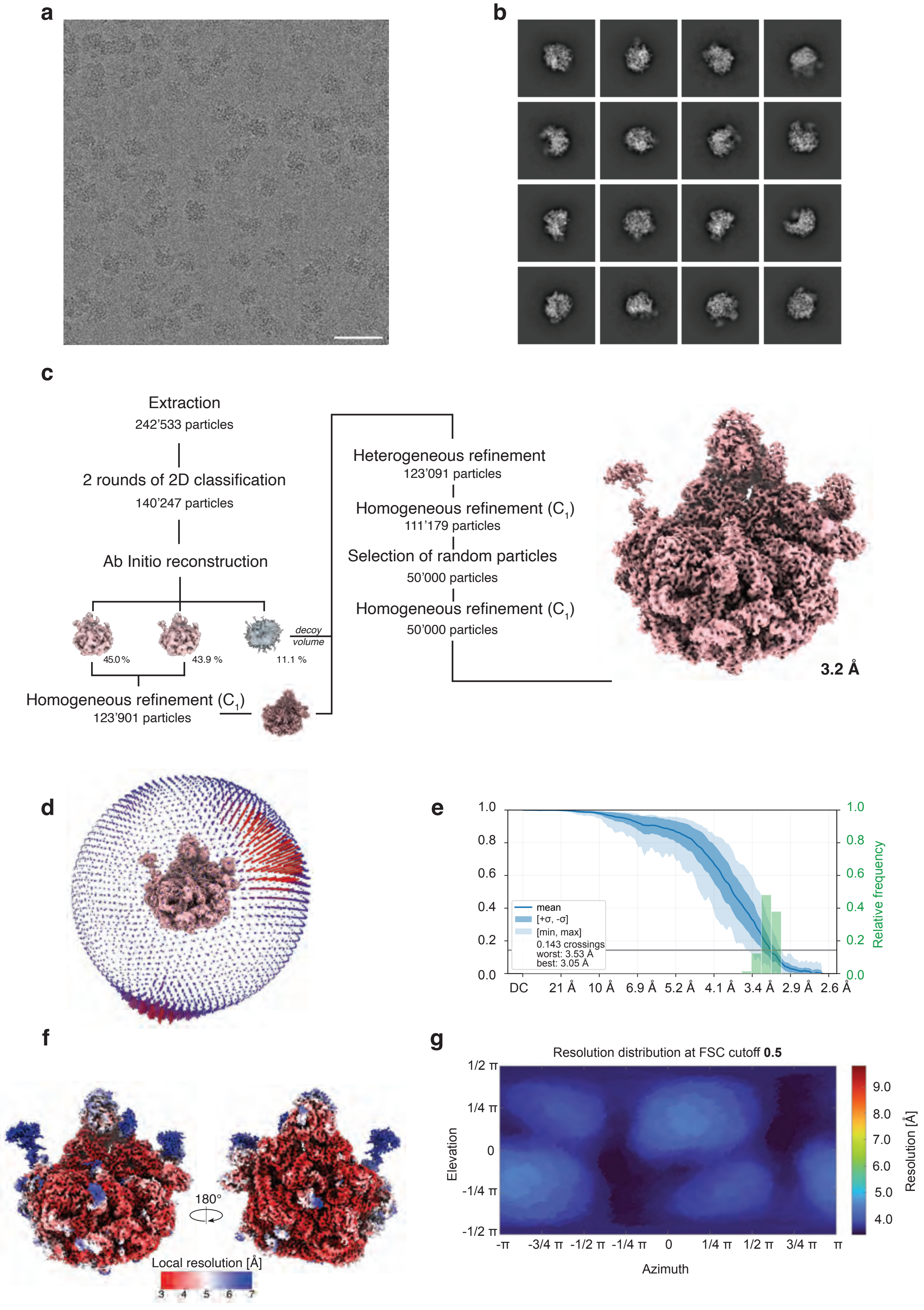
a
b
c
Extraction
242'533 particles
2 rounds of 2D classification
140'247 particles
Ab Initio reconstruction
decoy volume
45.0 %
43.9 %
11.1 %
Homogeneous refinement (C1)
123'901 particles
Heterogeneous refinement
123'091 particles
Homogeneous refinement (C1)
111'179 particles
Selection of random particles
50'000 particles
Homogeneous refinement (C1)
50'000 particles
3.2 Å
d
e
mean
[+σ, -σ]
[min, max]
0.143 crossings
worst: 3.53 Å
best: 3.05 Å
Relative frequency
DC 21 Å 10 Å 6.9 Å 5.2 Å 4.1 Å 3.4 Å 2.9 Å 2.6 Å
f
180°
Local resolution [Å]
3 4 5 6 7
g
Resolution distribution at FSC cutoff 0.5
Elevation
Azimuth
Resolution [Å]
-π -3/4 π -1/2 π -1/4 π 0 1/4 π 1/2 π 3/4 π π

**Supplementary Figure 5 | Cryo-EM data processing workflow for a revitrified sample of the 50S ribosome.** **a** Representative micrograph. Scale bar 100 Å. **b** 2D class averages **c** Data processing workflow in cryoSPARC. The symmetry applied is indicated in parentheses. A resolution of 3.2 Å was obtained in the final reconstruction. The map is shown at 2.8 $\sigma$. **d** Three-dimensional representation of angular distribution of the particles, with the color and height of the cylinders representing the frequency of observation (red: more abundant, blue: less abundant). **e** Plot of the 3DFSC with the 0.143 cutoff indicated (grey line). The mean FSC value is shown as a solid blue line, with dark blue shading corresponding to one standard deviation, and the light blue shading indicating the minimum and maximum values. The green bars represent a histogram of the resolution values obtained from the directional FSC. The more concise distribution of the minimum and maximum 0.143 crossing reflect the improvement of the preferred orientation of the 50S ribosomal subunit. **f** Final map with the local resolution estimation indicated in color. **g** The resolution distribution plot shown at FSC cutoff of 0.5, as obtained from the 3DFSC job in cryoSPARC. After revitrification, the resolution distribution also becomes more uniform.

a b

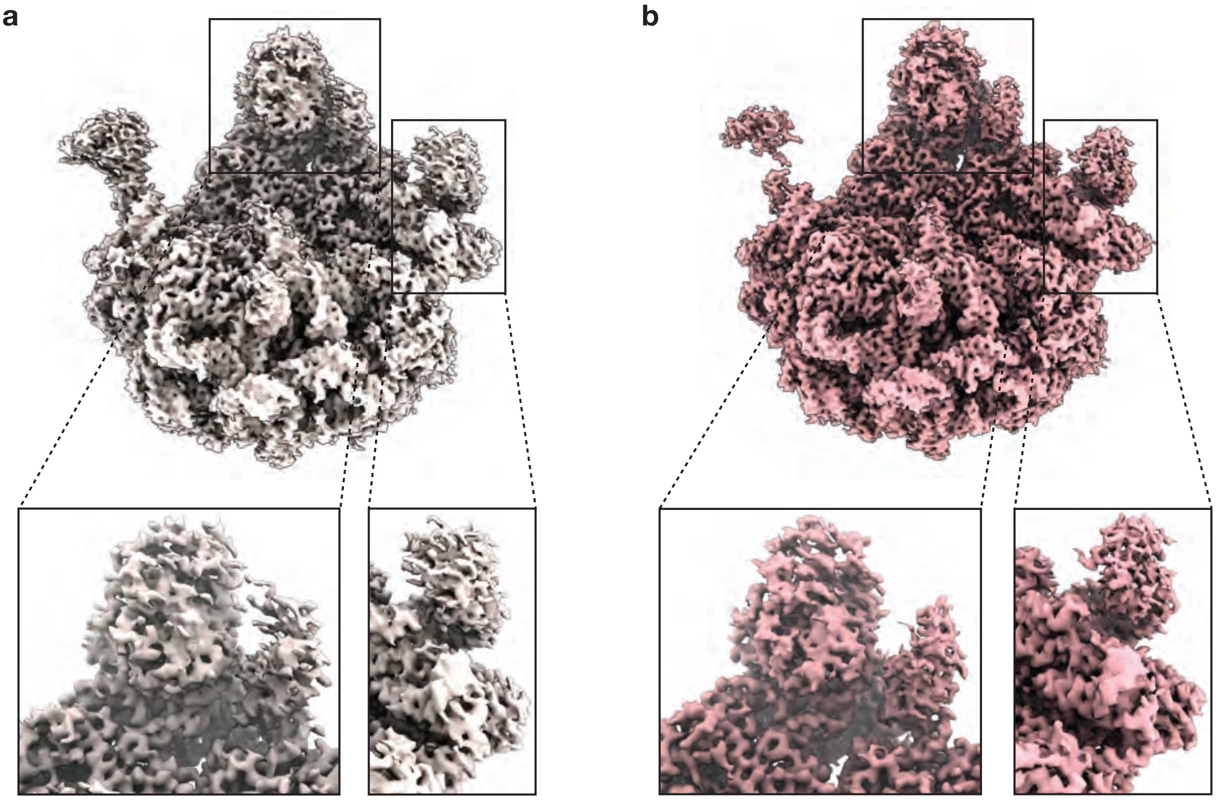

**Supplementary Figure 6 | Comparison of conventional and revitrified maps of the 50S ribosomal subunit. a** Conventional map. Insets highlight the streaky artefacts present in the map due to preferred orientation. **b** Revitrified map. Insets of same regions reveal the disappearance of the streaky artefacts thanks to a better angular sampling obtained after revitrification.

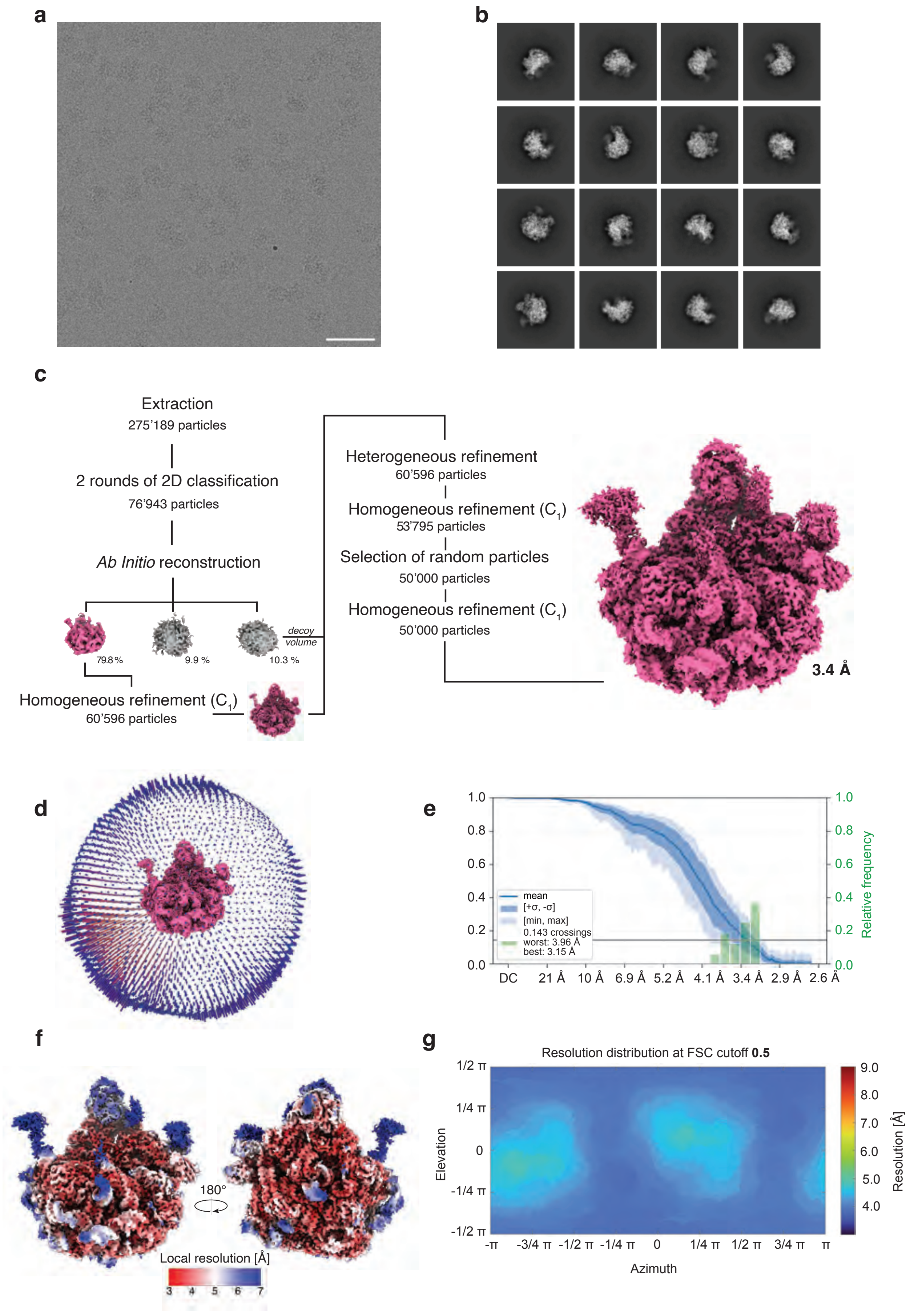
a
b
c
Extraction
275’189 particles
2 rounds of 2D classification
76’943 particles
Ab Initio reconstruction
79.8 %
9.9 %
10.3 %
decoy volume
Homogeneous refinement (C1)
60’596 particles
Heterogeneous refinement
60’596 particles
Homogeneous refinement (C1)
53’795 particles
Selection of random particles
50’000 particles
Homogeneous refinement (C1)
50’000 particles
3.4 Å
d
e
mean
[+σ, -σ]
[min, max]
0.143 crossings
worst: 3.96 Å
best: 3.15 Å
Relative frequency
DC 21 Å 10 Å 6.9 Å 5.2 Å 4.1 Å 3.4 Å 2.9 Å 2.6 Å
f
180°
Local resolution [Å]
3 4 5 6 7
g
Resolution distribution at FSC cutoff 0.5
Elevation
Azimuth
Resolution [Å]

**Supplementary Figure 7 | Cryo-EM data processing workflow for a deposited and revitrified sample of the 50S ribosome.** **a** Representative micrograph. Scale bar 100 Å. **b** 2D class averages **c** Data processing workflow in cryoSPARC. The symmetry applied is indicated in parentheses. A resolution of 3.4 Å was obtained in the final reconstruction. The map shown at 2.8 $\sigma$. **d** Three-dimensional representation of angular distribution of the particles, with the color and height of the cylinders representing the frequency of observation (red: more abundant, blue: less abundant). **e** Plot of the 3DFSC with the 0.143 cutoff indicated (grey line). The mean FSC value is shown as a solid blue line, with dark blue shading corresponding to one standard deviation, and the light blue shading indicating the minimum and maximum values. The green bars represent a histogram of the resolution values obtained from the directional FSC. The more concise distribution of the minimum and maximum 0.143 crossing reflect the improvement of the preferred orientation of the 50S ribosomal subunit. **f** Final map with the local resolution estimation indicated in color. **g** The resolution distribution plot shown at FSC cutoff of 0.5, as obtained from the 3DFSC job in cryoSPARC. Due to the increase in ice thickness after deposition, the directional resolutions, as well as the overall resolution of the reconstruction, decreased minimally compared to the reconstruction obtained from the revitrification-only experiment (Fig. S6g). Compared to the control, the directional resolution distribution improved (Fig. S5g).

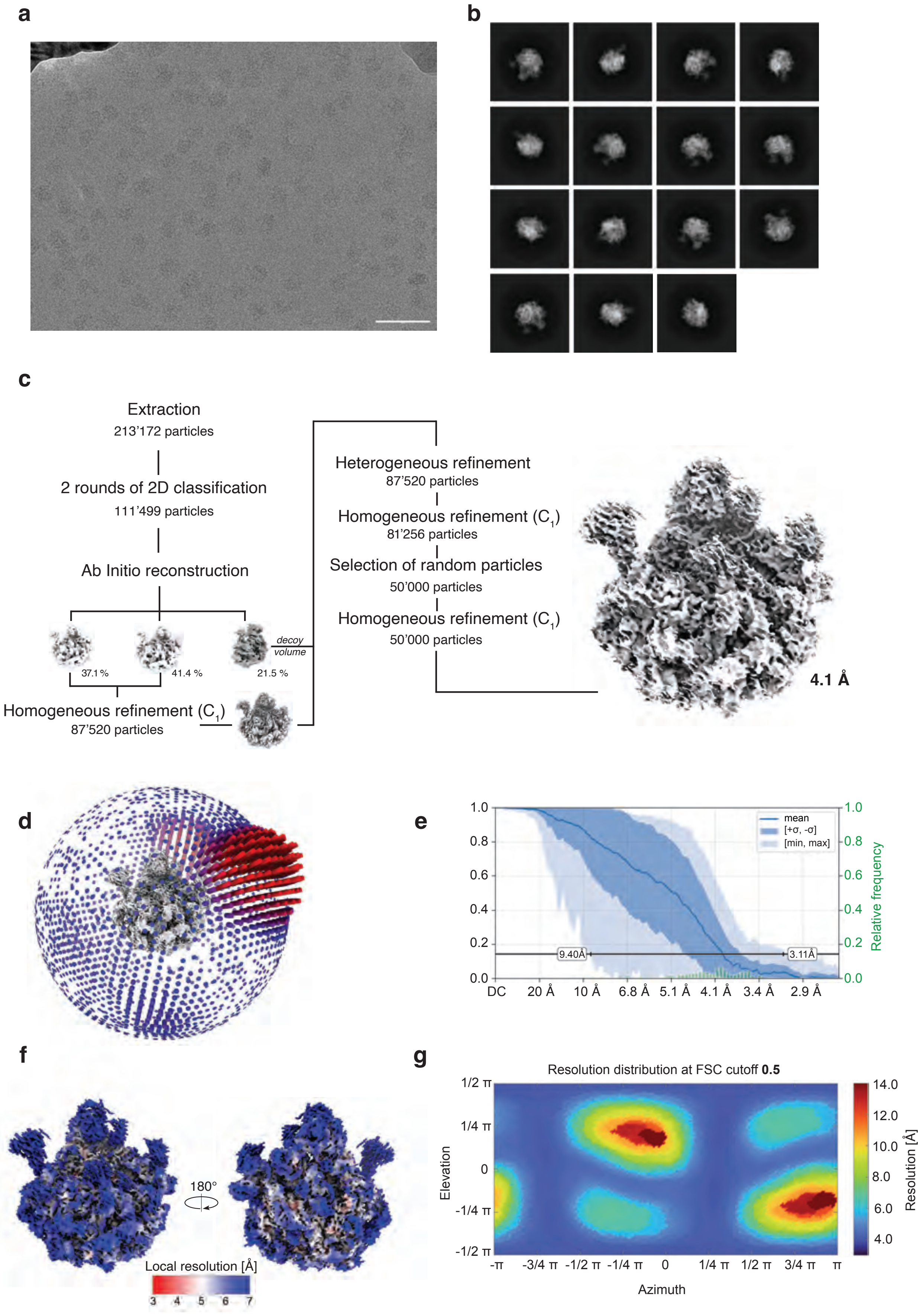

a
b
c
Extraction
213’172 particles
2 rounds of 2D classification
111’499 particles
Ab Initio reconstruction
decoy volume
37.1 %
41.4 %
21.5 %
Homogeneous refinement ($C_1$)
87’520 particles
Heterogeneous refinement
87’520 particles
Homogeneous refinement ($C_1$)
81’256 particles
Selection of random particles
50’000 particles
Homogeneous refinement ($C_1$)
50’000 particles
4.1 Å
d
e
mean
[+σ, -σ]
[min, max]
Relative frequency
9.40Å
3.11Å
DC 20 Å 10 Å 6.8 Å 5.1 Å 4.1 Å 3.4 Å 2.9 Å
f
180°
Local resolution [Å]
3 4 5 6 7
g
Resolution distribution at FSC cutoff **0.5**
Elevation
Azimuth
Resolution [Å]
-π -3/4 π -1/2 π -1/4 π 0 1/4 π 1/2 π 3/4 π π

**Supplementary Figure 8 | Cryo-EM data processing workflow for a conventional sample of the 50S ribosome, used as the control for shaped pulse revitrification. a** Representative micrograph. Scale bar 500 Å. **b** 2D class averages **c** Data processing workflow in cryoSPARC. The symmetry applied is indicated in parentheses. A resolution of 4.1 Å was obtained in the final reconstruction. The map is shown at 3 $\sigma$. **d** Three-dimensional representation of angular distribution of the particles, with the color and height of the cylinders representing the frequency of observation (red: more abundant, blue: less abundant). **e** Plot of the 3DFSC with the 0.143 cutoff indicated (grey line). The mean FSC value is shown as a solid blue line, with dark blue shading corresponding to one standard deviation, and the light blue shading indicating the minimum and maximum values. The green bars represent a histogram of the resolution values obtained from the directional FSC. The broad distribution of the minimum and maximum 0.143 crossing underline the preferred orientation of the 50S ribosomal subunit. **f** Final map with the local resolution estimation indicated in color. **g** The resolution distribution plot shown at FSC cutoff of 0.5, as obtained from the 3DFSC job in cryoSPARC. The plot shows regions with lower resolutions, corresponding to missing views in the angular distribution (panel d).

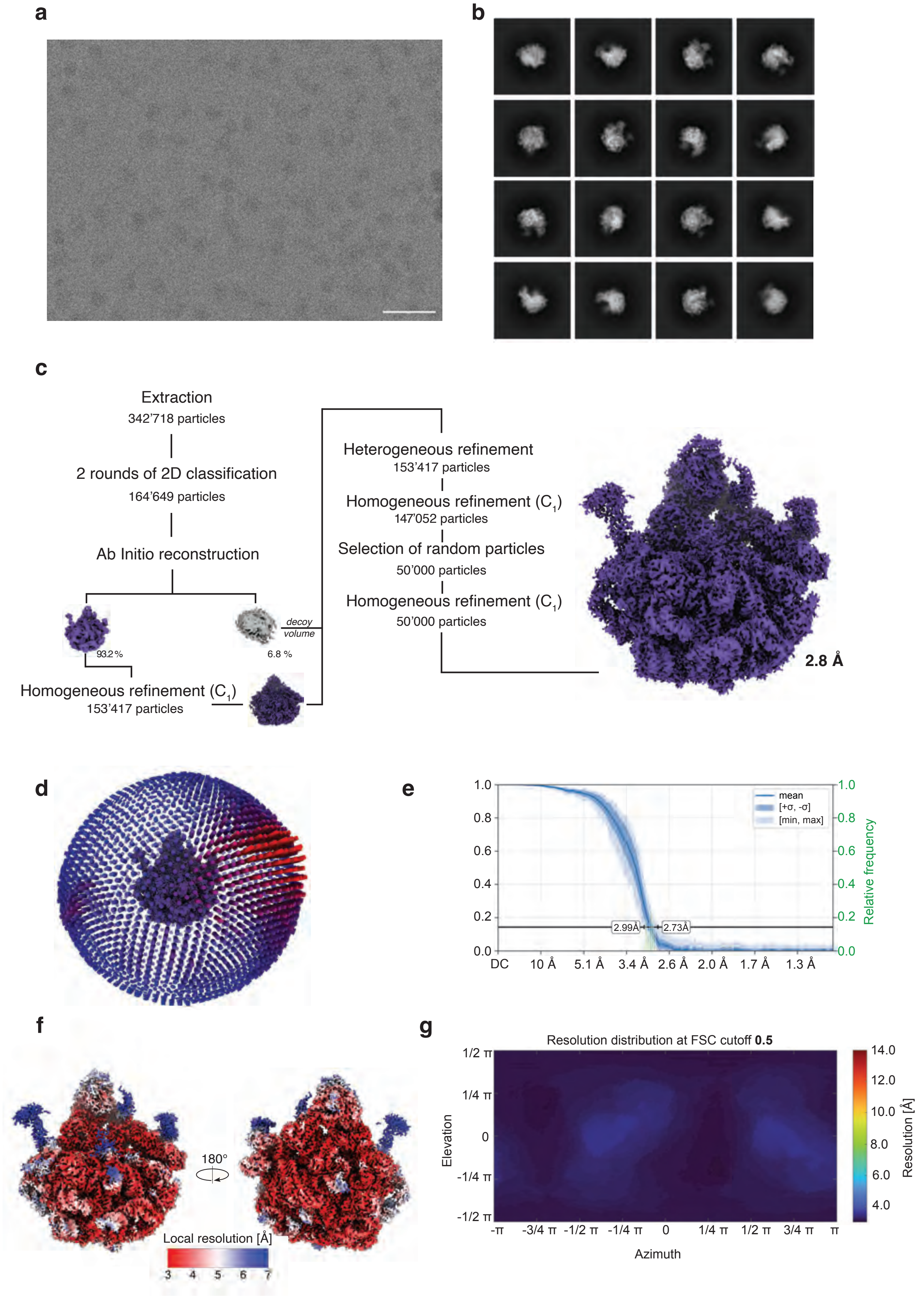
a
b
c
Extraction
342’718 particles
2 rounds of 2D classification
164’649 particles
Ab Initio reconstruction
decoy volume
93.2 %
6.8 %
Homogeneous refinement ($C_1$)
153’417 particles
Heterogeneous refinement
153’417 particles
Homogeneous refinement ($C_1$)
147’052 particles
Selection of random particles
50’000 particles
Homogeneous refinement ($C_1$)
50’000 particles
2.8 Å
d
e
mean
[+σ, -σ]
[min, max]
Relative frequency
2.99Å
2.73Å
DC
10 Å
5.1 Å
3.4 Å
2.6 Å
2.0 Å
1.7 Å
1.3 Å
f
180°
Local resolution [Å]
3
4
5
6
7
g
Resolution distribution at FSC cutoff **0.5**
Elevation
Azimuth
Resolution [Å]

**Supplementary Figure 9 | Cryo-EM data processing workflow for a shaped pulse revitrified sample of the 50S ribosome. a** Representative micrograph. Scale bar 500 Å. **b** 2D class averages **c** Data processing workflow in cryoSPARC. The symmetry applied is indicated in parentheses. A resolution of 2.9 Å was obtained in the final reconstruction. The map is shown at 3 $\sigma$. **d** Three-dimensional representation of angular distribution of the particles, with the color and height of the cylinders representing the frequency of observation (red: more abundant, blue: less abundant). **e** Plot of the 3DFSC with the 0.143 cutoff indicated (grey line). The mean FSC value is shown as a solid blue line, with dark blue shading corresponding to one standard deviation, and the light blue shading indicating the minimum and maximum values. The green bars represent a histogram of the resolution values obtained from the directional FSC. The narrow distribution of the minimum and maximum 0.143 crossing reflect the improvement of the preferred orientation of the 50S ribosomal subunit. **f** Final map with the local resolution estimation indicated in color. **g** The resolution distribution plot shown at FSC cutoff of 0.5, as obtained from the 3DFSC job in cryoSPARC. Compared to the control, the directional resolution distribution improved significantly (Fig. S8g).

**3 | Data collection and analysis – HIV-1 Envelope ectodomain protein**

The HIV-1 datasets were collected on a Titan Krios G4 using a Falcon 4i camera at the Dubochet Center for Imaging in Lausanne, Switzerland. The movies were recorded in eer format using a magnification of 96’000 x, resulting in a pixel size of 0.83 Å per pixel. A total dose of 50 $e^-/Å^2$ was used and the defocus range was set to -1 µm to -2.4 µm.

For the conventional HIV-1 dataset, 2’614 micrographs were collected. For the revitrified dataset, data was collected from two different grids; from the first grid, 5’447 micrographs were collected, whereas from the second grid 5’081 images were recorded. All the datasets were subjected patch motion correction, patch CTF estimation and a manual curation based on CTF resolution estimation, relative ice thickness and the defocus range. Next, the particles were picked using a blob picker with a diameter of 100 – 150 Å. The particles were extracted with a box size of 480 px and down sampled to 240 px. The particles from the two revitrified datasets were combined before sorting in 2 rounds of 2D classification. The selected particles were subjected to an *ab initio* reconstruction using two classes. Particles were further classified in 3D through two rounds of heterogeneous refinements, where all selected particles from the first round of 2D classification were sorted against the good initial volume and five decoy classes, generated through the pre-mature termination of an *ab initio* reconstruction. The selected set of particles was then refined in a non-uniform refinement with $C_1$ symmetry. Next, the particles were re-extracted at full box size, locally and globally CTF-refined before being directed into another non-uniform refinement, without dynamic masking and symmetry enforcement. Finally, 50’000 random particles were selected and again refined in a non-uniform refinement without dynamic masking using $C_1$ symmetry and the filtered conventional volume as a reference, before being directed into an orientation diagnostics job.

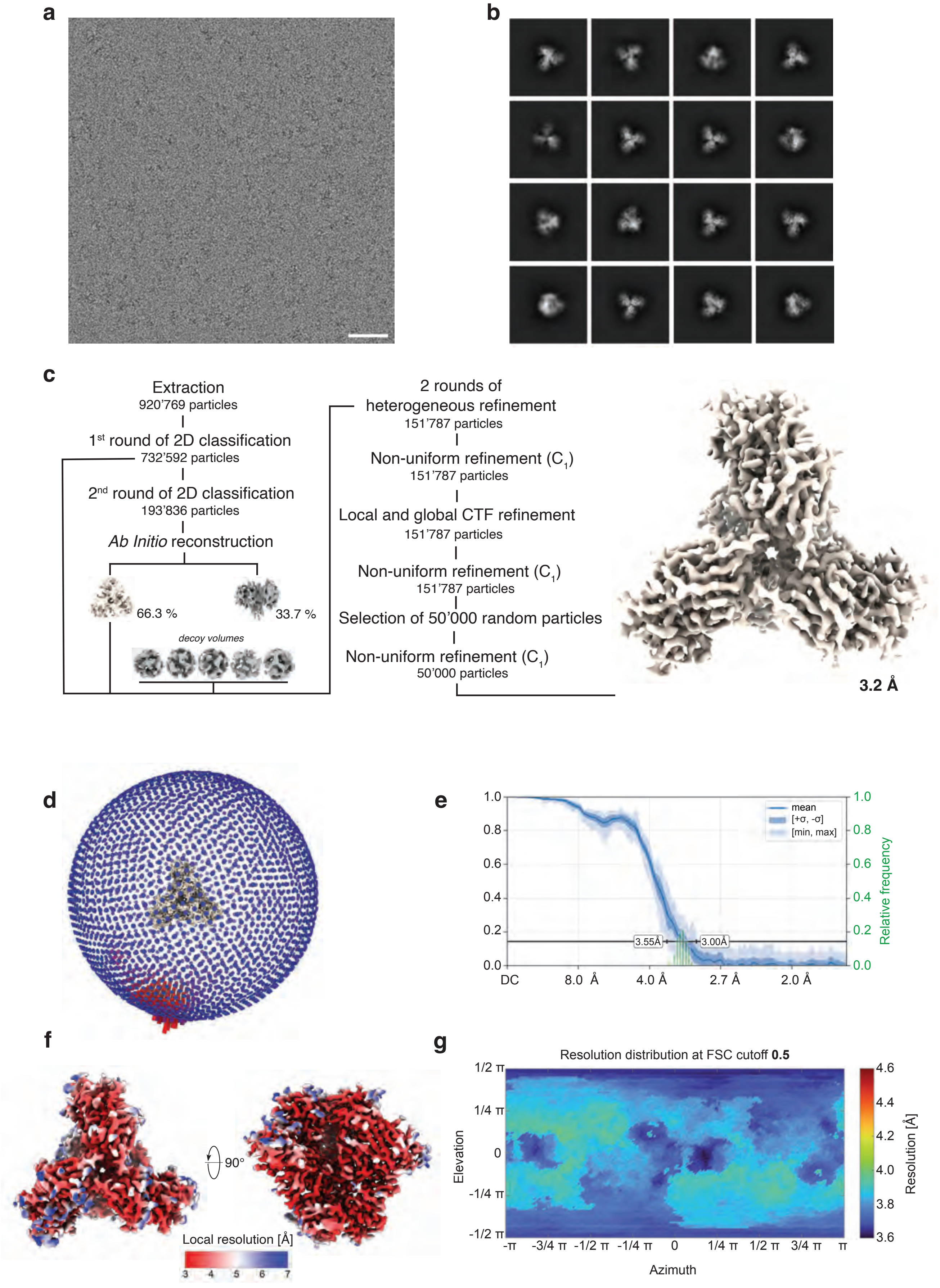

a
b
c
Extraction
920’769 particles
1st round of 2D classification
732’592 particles
2nd round of 2D classification
193’836 particles
Ab Initio reconstruction
66.3 %
33.7 %
decoy volumes
2 rounds of heterogeneous refinement
151’787 particles
Non-uniform refinement (C1)
151’787 particles
Local and global CTF refinement
151’787 particles
Non-uniform refinement (C1)
151’787 particles
Selection of 50’000 random particles
Non-uniform refinement (C1)
50’000 particles
3.2 Å
d
e
mean
[+σ, -σ]
[min, max]
Relative frequency
3.55Å
3.00Å
DC
8.0 Å
4.0 Å
2.7 Å
2.0 Å
f
90°
Local resolution [Å]
g
Resolution distribution at FSC cutoff 0.5
Elevation
Azimuth
Resolution [Å]

**Supplementary Figure 10 | Cryo-EM data processing workflow for a conventional sample of the HIV-1 Envelope ectodomain protein. a** Representative micrograph. Scale bar 100 Å. **b** 2D class averages. **c** Data processing workflow in cryoSPARC. The symmetry applied is indicated in parentheses. A resolution of 3.2 Å was obtained in the final reconstruction. The map is shown at 15 $\sigma$. **d** Three-dimensional representation of angular distribution of the particles, with the color and height of the cylinders representing the frequency of observation (red: more abundant, blue: less abundant). **e** Plot of the 3DFSC with the 0.143 cutoff indicated (grey line). The mean FSC value is shown as a solid blue line, with dark blue shading corresponding to one standard deviation, and the light blue shading indicating the minimum and maximum values. The green bars represent a histogram of the resolution values obtained from the directional FSC. **f** Final map with the local resolution estimation indicated in color. **g** The resolution distribution plot shown at FSC cutoff of 0.5, as obtained from the 3DFSC job in cryoSPARC.

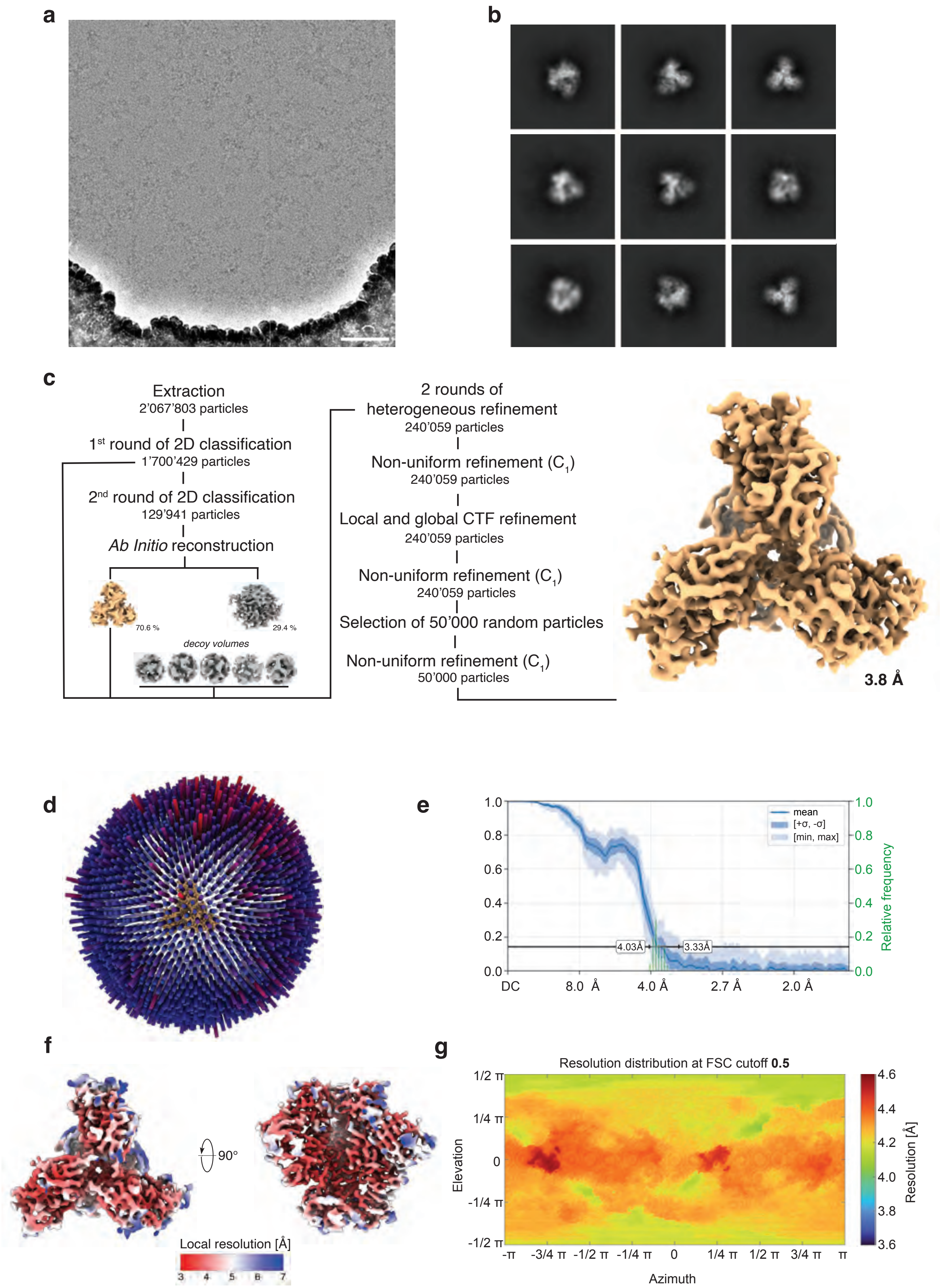
a
b
c
Extraction
2'067'803 particles
1st round of 2D classification
1'700'429 particles
2nd round of 2D classification
129'941 particles
Ab Initio reconstruction
70.6 %
29.4 %
decoy volumes
2 rounds of
heterogeneous refinement
240'059 particles
Non-uniform refinement (C1)
240'059 particles
Local and global CTF refinement
240'059 particles
Non-uniform refinement (C1)
240'059 particles
Selection of 50'000 random particles
Non-uniform refinement (C1)
50'000 particles
3.8 Å
d
e
mean
[+σ, -σ]
[min, max]
Relative frequency
4.03Å
3.33Å
DC
8.0 Å
4.0 Å
2.7 Å
2.0 Å
f
90°
Local resolution [Å]
3
4
5
6
7
g
Resolution distribution at FSC cutoff 0.5
Elevation
Azimuth
Resolution [Å]

**Supplementary Figure 11 | Cryo-EM data processing workflow for a revitrified sample of the HIV-1 Envelope ectodomain protein. a** Representative micrograph. Scale bar 100 Å. **b** 2D class averages. **c** Data processing workflow in cryoSPARC. The symmetry applied is indicated in parentheses. A resolution of 3.8 Å was obtained in the final reconstruction. The map is contoured at 15 $\sigma$. **d** Three-dimensional representation of angular distribution of the particles, with the color and height of the cylinders representing the frequency of observation (red: more abundant, blue: less abundant). **e** Plot of the 3DFSC with the 0.143 cutoff indicated (grey line). The mean FSC value is shown as a solid blue line, with dark blue shading corresponding to one standard deviation, and the light blue shading indicating the minimum and maximum values. The green bars represent a histogram of the resolution values obtained from the directional FSC. **f** Final map with the local resolution estimation indicated in color. **g** The resolution distribution plot shown at FSC cutoff of 0.5, as obtained from the 3DFSC job in cryoSPARC. Despite the improvement in angular distribution, the resolution decreased in the revitrified sample, which we attributed to the presence of astigmatism in the microscope during data collection.

### 4 | Data collection and analysis – Hemagglutinin

The four Hemagglutinin datasets were recorded on a Titan Krios G4 equipped with a Falcon 4i and a SelectrisX energy filter at the Dubochet Center for Imaging in Lausanne, Switzerland. The movies were collected in eer format at a magnification of 165'000 x, corresponding to a pixel size of 0.732 Å. A total dose of 40 $e^{-}/Å^{2}$ was used for the conventional, revitrified, and leading-edge revitrified and deposited and revitrified dataset, respectively. A defocus range of -0.8 µm to -2.5 µm and an energy filter slit of 10 eV was applied for all four datasets.

For the conventional, revitrified, deposited and revitrified and shaped pulse revitrified HA samples, 2'251, 6'476, 10'114, and 10'270 micrographs were collected. The micrographs were subjected to patch motion correction and patch CTF estimation. The micrographs were manually curated based on their CTF resolution estimation, relative ice thickness and total full-frame motion, resulting in 1'651, 4'608, 7'982, and 8'590 high-quality micrographs. On 100 micrographs of each dataset, a denoiser was trained and subsequently applied to the curated micrographs. Particles were picked from denoised micrographs using a blob picker with a diameter of 80 – 140 Å. Next, the particles were extracted from the raw, non-denoised micrographs using a box size of 360 px, which was cropped to 180 px. The extracted particles were subjected to one to two rounds of 2D classification, followed by an *ab initio* reconstruction with two classes. The particles were then further cleaned up in one round of heterogeneous refinement using a decoy volume and the best resolved *ab initio* volume as inputs. The sorted particles were then refined in a homogeneous refinement, before being re-extracted at full box size. Out of these, 50'000 random particles were selected and then subjected into a final homogeneous refinement with $C_3$ symmetry, using the filtered conventional map as a reference, and an orientation diagnostics job.

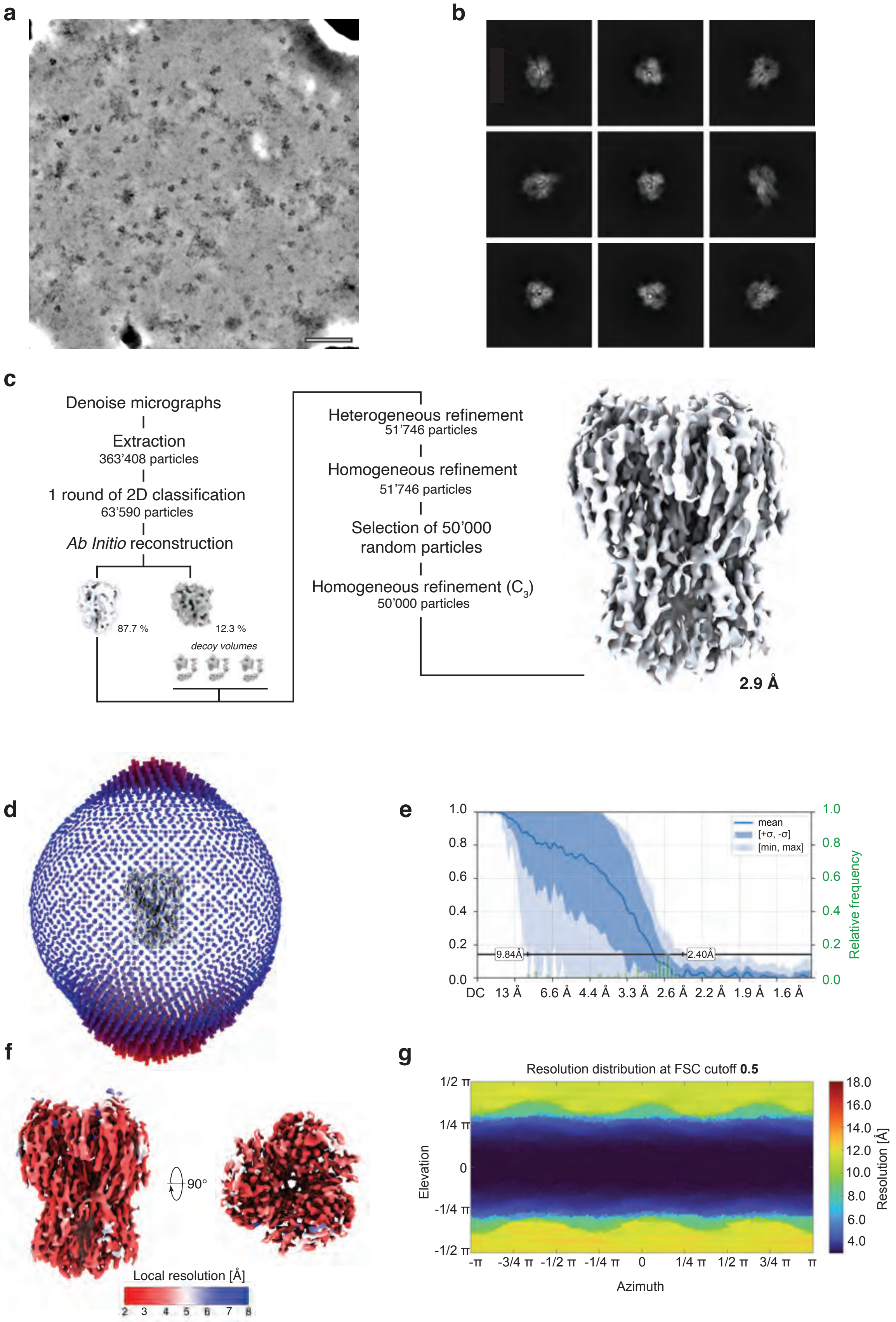
a
b
c
Denoise micrographs
Extraction
363'408 particles
1 round of 2D classification
63'590 particles
Ab Initio reconstruction
87.7 %
12.3 %
decoy volumes
Heterogeneous refinement
51'746 particles
Homogeneous refinement
51'746 particles
Selection of 50'000
random particles
Homogeneous refinement ($C_3$)
50'000 particles
2.9 Å
d
e
mean
[+σ, -σ]
[min, max]
Relative frequency
9.84Å
2.40Å
DC 13 Å 6.6 Å 4.4 Å 3.3 Å 2.6 Å 2.2 Å 1.9 Å 1.6 Å
f
90°
Local resolution [Å]
2 3 4 5 6 7 8
g
Resolution distribution at FSC cutoff **0.5**
Elevation
Azimuth
Resolution [Å]
-π -3/4 π -1/2 π -1/4 π 0 1/4 π 1/2 π 3/4 π π
1/2 π 1/4 π 0 -1/4 π -1/2 π

**Supplementary Figure 12 | Cryo-EM data processing workflow for a conventional sample of Hemagglutinin. a** Representative denoised micrograph. Scale bar 100 Å. **b** 2D class averages. **c** Data processing workflow in cryoSPARC. The final map reconstructed to 2.9 Å (as reported by cryoSPARC. The apparent resolution, due to anisotropy is worse). The symmetry applied is indicated in parentheses. The map is shown 7 $\sigma$. **d** Three-dimensional representation of angular distribution of the particles, with the color and height of the cylinders representing the frequency of observation (red: more abundant, blue: less abundant). **e** Plot of the 3DFSC with the 0.143 cutoff indicated (grey line). The mean FSC value is shown as a solid blue line, with dark blue shading corresponding to one standard deviation, and the light blue shading indicating the minimum and maximum values. The green bars represent a histogram of the resolution values obtained from the directional FSC. **f** Final map with the local resolution estimation indicated in color. **g** The resolution distribution plot shown at FSC cutoff of 0.5, as obtained from the 3DFSC job in cryoSPARC.

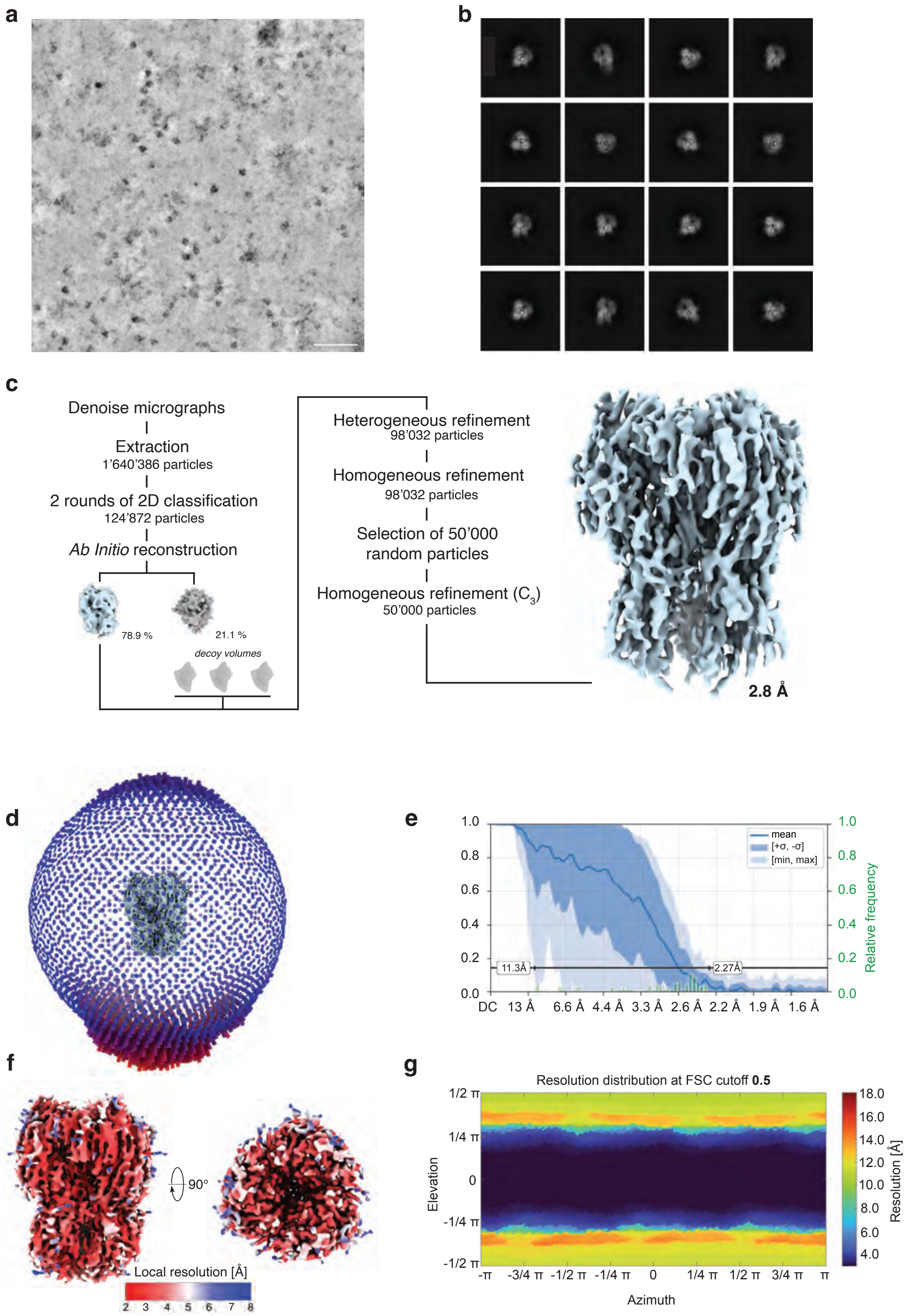
a
b
c
Denoise micrographs
Extraction
1’640’386 particles
2 rounds of 2D classification
124’872 particles
Ab Initio reconstruction
78.9 %
21.1 %
decoy volumes
Heterogeneous refinement
98’032 particles
Homogeneous refinement
98’032 particles
Selection of 50’000
random particles
Homogeneous refinement (C3)
50’000 particles
2.8 Å
d
e
mean
[+σ, -σ]
[min, max]
11.3Å
2.27Å
DC 13 Å 6.6 Å 4.4 Å 3.3 Å 2.6 Å 2.2 Å 1.9 Å 1.6 Å
Relative frequency
f
90°
Local resolution [Å]
2 3 4 5 6 7 8
g
Resolution distribution at FSC cutoff 0.5
Elevation
Azimuth
Resolution [Å]

**Supplementary Figure 13 | Cryo-EM data processing workflow for a revitrified sample of Hemagglutinin.** **a** Representative denoised micrograph. Scale bar 100 Å. **b** 2D class averages **c** Data processing workflow in cryoSPARC. The final map reconstructed to 2.8 Å (as reported by cryoSPARC. The apparent resolution, due to anisotropy is worse). The symmetry applied is indicated in parentheses. The map is shown at 7 $\sigma$. **d** Three-dimensional representation of angular distribution of the particles, with the color and height of the cylinders representing the frequency of observation (red: more abundant, blue: less abundant). **e** Plot of the 3DFSC with the 0.143 cutoff indicated (grey line). The mean FSC value is shown as a solid blue line, with dark blue shading corresponding to one standard deviation, and the light blue shading indicating the minimum and maximum values. The green bars represent a histogram of the resolution values obtained from the directional FSC. **f** Final map with the local resolution estimation indicated in color. **g** The resolution distribution plot shown at FSC cutoff of 0.5, as obtained from the 3DFSC job in cryoSPARC.

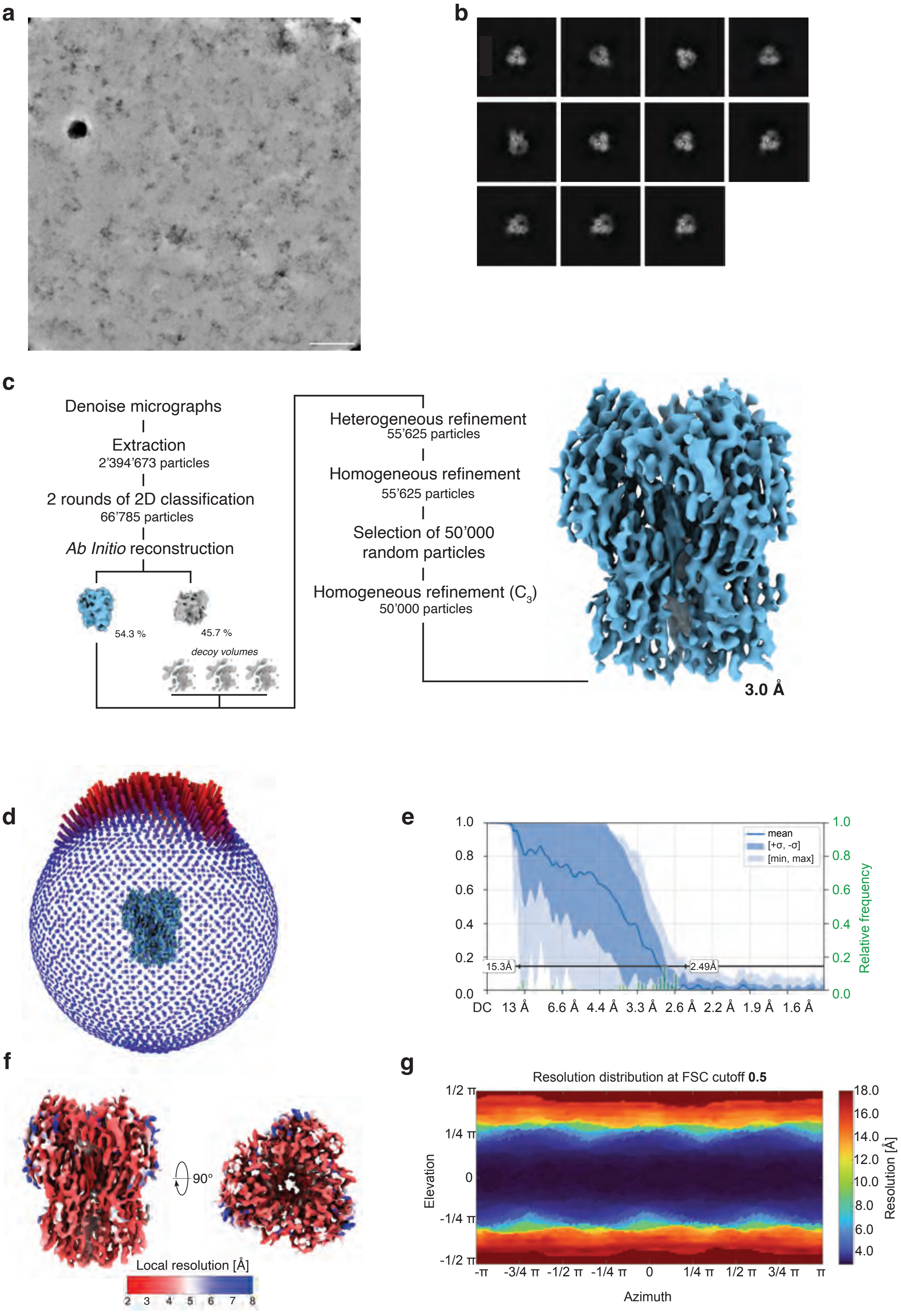
a
b
c
Denoise micrographs
Extraction
2'394'673 particles
2 rounds of 2D classification
66'785 particles
Ab Initio reconstruction
54.3 %
45.7 %
decoy volumes
Heterogeneous refinement
55'625 particles
Homogeneous refinement
55'625 particles
Selection of 50'000
random particles
Homogeneous refinement (C3)
50'000 particles
3.0 Å
d
e
1.0
0.8
0.6
0.4
0.2
0.0
mean
[+σ, -σ]
[min, max]
15.3Å
2.49Å
Relative frequency
DC 13 Å 6.6 Å 4.4 Å 3.3 Å 2.6 Å 2.2 Å 1.9 Å 1.6 Å
f
90°
Local resolution [Å]
2 3 4 5 6 7 8
g
Resolution distribution at FSC cutoff 0.5
1/2 π
1/4 π
0
-1/4 π
-1/2 π
Elevation
-π -3/4 π -1/2 π -1/4 π 0 1/4 π 1/2 π 3/4 π π
Azimuth
18.0
16.0
14.0
12.0
10.0
8.0
6.0
4.0
Resolution [Å]

**Supplementary Figure 14 | Cryo-EM data processing workflow for a deposited and revitrified sample of Hemagglutinin.** **a** Representative denoised micrograph. Scale bar 100 Å. **b** 2D class averages. **c** Data processing workflow in cryoSPARC. The final map reconstructed to 3.0 Å (as reported by cryoSPARC. The apparent resolution, due to anisotropy is worse). The symmetry applied is indicated in parentheses. The map is shown at 7 $\sigma$. **d** Three-dimensional representation of angular distribution of the particles, with the color and height of the cylinders representing the frequency of observation (red: more abundant, blue: less abundant). **e** Plot of the 3DFSC with the 0.143 cutoff indicated (grey line). The mean FSC value is shown as a solid blue line, with dark blue shading corresponding to one standard deviation, and the light blue shading indicating the minimum and maximum values. The green bars represent a histogram of the resolution values obtained from the directional FSC. **f** Final map with the local resolution estimation indicated in color. **g** The resolution distribution plot shown at FSC cutoff of 0.5, as obtained from the 3DFSC job in cryoSPARC. The increase in ice thickness after deposition leads to a decrease in the resolution distribution.

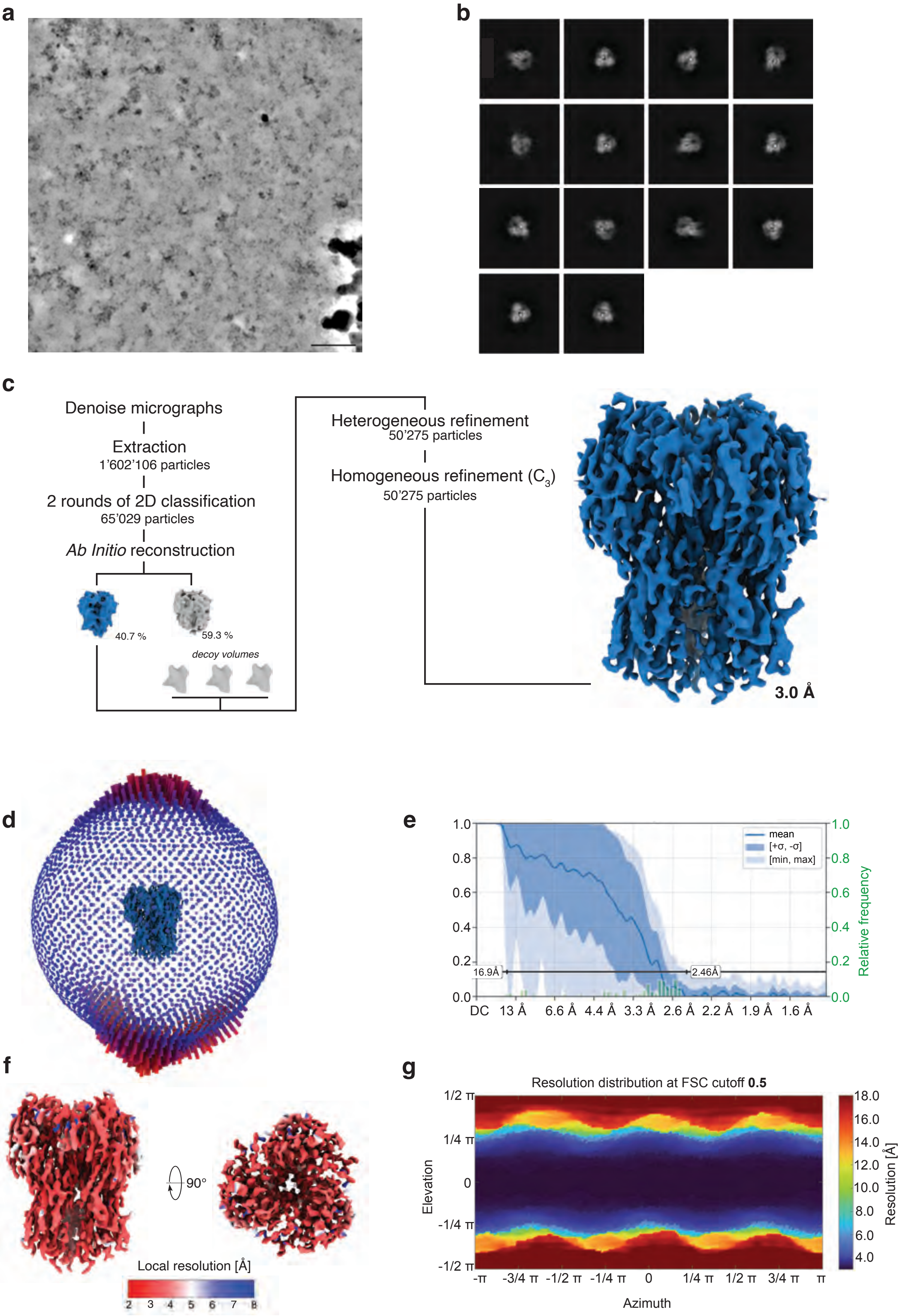
a
b
c
Denoise micrographs
Extraction
1'602'106 particles
2 rounds of 2D classification
65'029 particles
Ab Initio reconstruction
40.7 %
59.3 %
decoy volumes
Heterogeneous refinement
50'275 particles
Homogeneous refinement (C3)
50'275 particles
3.0 Å
d
e
mean
[+σ, -σ]
[min, max]
Relative frequency
16.9Å
2.46Å
DC 13 Å 6.6 Å 4.4 Å 3.3 Å 2.6 Å 2.2 Å 1.9 Å 1.6 Å
f
90°
Local resolution [Å]
2 3 4 5 6 7 8
g
Resolution distribution at FSC cutoff 0.5
Elevation
Azimuth
Resolution [Å]

**Supplementary Figure 15 | Cryo-EM data processing workflow for a shaped pulse revitrified sample of Hemagglutinin.** **a** Representative denoised micrograph. Scale bar 100 Å. **b** 2D class averages. **c** Data processing workflow in cryoSPARC. The final map reconstructed to 3.0 Å (as reported by cryoSPARC. The apparent resolution, due to anisotropy is worse). The symmetry applied is indicated in parentheses. The map is shown at 7 $\sigma$. **d** Three-dimensional representation of angular distribution of the particles, with the color and height of the cylinders representing the frequency of observation (red: more abundant, blue: less abundant). **e** Plot of the 3DFSC with the 0.143 cutoff indicated (grey line). The mean FSC value is shown as a solid blue line, with dark blue shading corresponding to one standard deviation, and the light blue shading indicating the minimum and maximum values. The green bars represent a histogram of the resolution values obtained from the directional FSC. **f** Final map with the local resolution estimation indicated in color. **g** The resolution distribution plot shown at FSC cutoff of 0.5, as obtained from the 3DFSC job in cryoSPARC.

### 5 | Simulation of the temperature evolution of the sample

The temperature evolution of the sample under laser irradiation was simulated with COMSOL Multiphysics 6.1, as previously described [2,3]. Figure S16a schematically displays the shape of a rectangular pulse as used in the experiments of Figs. 1 and 3 together with the resulting temperature evolution of the sample. We report the average sample temperature within a hole in the gold film in the center of the laser spot. Here, a laser power of 150 mW was simulated, for which the sample temperature reaches 285 K at the end of the laser pulse. For comparison, Fig. S16b shows the simulated temperature evolution of the sample for a shaped pulse with an intense leading edge as used in the experiments of Fig. 2 (1 μs duration of the initial spike with 10 times the laser power). The simulation revealed that the sample temperature rises more rapidly at the beginning of the laser pulse and overshoots the plateau temperature for a duration of approximately 1.5 μs, reaching a maximum of approximately 310 K. Note that while schematic laser pulse shapes are shown in Fig. S16, experimentally determined pulse shapes were used in the simulations as recorded with a fast photodiode.

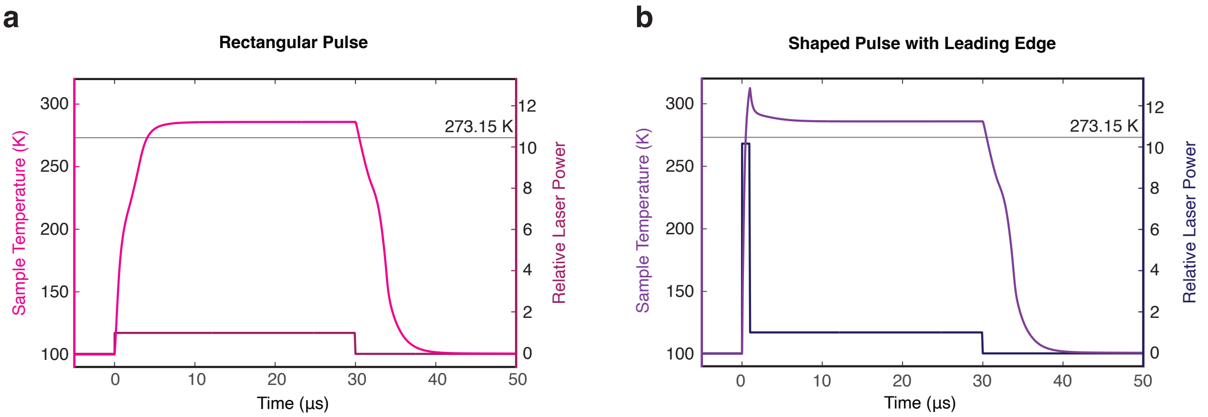


**Supplementary Figure 16 | Simulation of the sample temperature evolution under irradiation with a rectangular and a shaped laser pulse. a** Simulated temperature evolution of the sample (pink) under irradiation with a rectangular laser pulse (schematically shown in magenta). **b** Under irradiation with a shaped laser pulse (navy, 1 μs initial spike of 10 times the power), the sample heats up more rapidly, and the temperature briefly overshoots (shown in purple).

### 6 | Cryo-EM data collection, refinement and validation statistics

**Supplementary Table 1 |** Cryo-EM statistics for the datasets collected on the T20S proteasome

| | T20S conventional | T20S revitrified | T20S revitrified after deposition |
|---|---|---|---|
| | EMDB-51744<br>EMPIAR-12389 | EMDB-51745<br>EMPIAR-12388 | EMDB-51746<br>EMPIAR-12390 |
| **Data collection and processing** | | | |
| Microscope | Titan Krios G3i | Titan Krios G3i | Titan Krios G3i |
| Camera | K3 | K3 | K3 |
| Energy Filter (slit) | BioQuantum (20 eV) | BioQuantum (20 eV) | BioQuantum (20 eV) |
| Magnification | 130'000 | 130'000 | 130'000 |
| Voltage (kV) | 300 | 300 | 300 |
| Electron exposure ($e^-/Å^2$) | 67.7 | 65.3 | 67.7 |
| Defocus range (µm) | -0.6 to -2.0 | -0.6 to -2.0 | -0.6 to -2.0 |
| Pixel size (Å) | 0.651 | 0.651 | 0.651 |
| Symmetry imposed | $D_7$ | $D_7$ | $D_7$ |
| Initial particle images (no.) | 397’944 | 450’862 | 409’644 |
| Final particle images (no.) | 50’000 | 50’000 | 50’000 |
| Map resolution (Å) | 2.6 | 2.5 | 2.8 |
| FSC threshold | 0.143 | 0.143 | 0.143 |
| Map sharpening *B* factor ($Å^2$) | 82.7 | 79.7 | 95.0 |
| Orientation Diagnostics | | | |
| cFAR | 0.82 | 0.85 | 0.83 |
| SCF* | 0.88 | 0.95 | 0.97 |

**Supplementary Table 2 |** Cryo-EM statistics for the datasets collected on the 50S ribosomal subunit

| | 50S conventional<br><br>EMDB-51747<br>EMPIAR-12397 | 50S revitrified<br><br>EMDB-51748<br>EMPIAR-12398 | 50S revitrified after deposition<br><br>EMDB-51749<br>EMPIAR-12399 | 50S shaped pulse conventional<br><br>EMDB-51750<br>EMPIAR-12435 | 50S shaped pulse revitrified<br><br>EMDB-51751<br>EMPIAR-12436 |
|---|---|---|---|---|---|
| **Data collection and processing** | | | | | |
| Microscope | Titan Krios G4 | Titan Krios G4 | Titan Krios G4 | Titan Krios G3 | Titan Krios G3 |
| Camera | Falcon 4i | Falcon 4i | Falcon 4i | K3 | K3 |
| Energy Filter (slit) | none | none | none | BioQuantum (20 eV) | BioQuantum (20 eV) |
| Magnification | 120'000 | 120'000 | 120'000 | 130'000 | 130'000 |
| Voltage (keV) | 300 | 300 | 300 | 300 | 300 |
| Electron exposure ($e^{-}$/Å$^2$) | 38.6 | 40.0 | 40.0 | 61.8 | 63.4 |
| Defocus range (μm) | -0.4 to -1.0 | -0.4 to -1.0 | -0.4 to -1.0 | -0.4 to -1.6 | -0.4 to -1.6 |
| Pixel size (Å) | 0.658 | 0.658 | 0.658 | 0.651 | 0.651 |
| Symmetry imposed | $C_1$ | $C_1$ | $C_1$ | $C_1$ | $C_1$ |
| Initial particle images (no.) | 296'144 | 242'533 | 275'189 | 373'837 | 342'718 |
| Final particle images (no.) | 50'000 | 50'000 | 50'000 | 50'000 | 50'000 |
| Map resolution (Å) | 3.5 | 3.2 | 3.4 | 4.1 | 2.9 |
| FSC threshold | 0.143 | 0.143 | 0.143 | 0.143 | 0.143 |
| Map sharpening *B* factor (Å$^2$) | 43.6 | 47.9 | 37.7 | 57.8 | 40.4 |
| Orientation Diagnostics | | | | | |
| cFAR | 0.10 | 0.51 | 0.50 | 0.03 | 0.70 |
| SCF* | 0.42 | 0.74 | 0.90 | 0.18 | 0.90 |

**Supplementary Table 3 |** Cryo-EM statistics for the datasets collected on the HIV-1 Envelope ectodomain protein

| | HIV-1 conventional<br><br>EMDB-51752<br>EMPIAR-12437 | HIV-1 revitrified<br><br>EMDB-51753<br>EMPIAR-12438 |
|---|---|---|
| **Data collection and processing** | | |
| Microscope | Titan Krios G4 | Titan Krios G4 |
| Camera | Falcon 4i | Falcon 4i |
| Energy Filter | none | none |
| Magnification | 96'000 | 96'000 |
| Voltage (kV) | 300 | 300 |
| Electron exposure ($e^{-}/Å^2$) | 50.0 | 50.0 |
| Defocus range (μm) | -1.0 to -2.4 | -1.0 to -2.4 |
| Pixel size (Å) | 0.830 | 0.830 |
| Symmetry imposed | $C_1$ | $C_1$ |
| Initial particle images (no.) | 920’769 | 2’067’803 |
| Final particle images (no.) | 50’000 | 50’000 |
| Map resolution (Å) | 3.2 | 3.8 |
| FSC threshold | 0.143 | 0.143 |
| Map sharpening *B* factor ($Å^2$) | 21.1 | 29.7 |
| Orientation Diagnostics | | |
| cFAR | 0.78 | 0.71 |
| SCF* | 0.89 | 0.99 |

**Supplementary Table 4 |** Cryo-EM Statistics for the datasets collected on Hemagglutinin (HA)

| | HA conventional | HA revitrified | HA revitrified after deposition | HA shaped pulse revitrified |
|---|---|---|---|---|
| | EMDB-51754<br>EMPIAR-12439 | EMDB-51755<br>EMPIAR-12440 | EMDB-51756<br>EMPIAR-12441 | EMDB-51757<br>EMPIAR-12442 |
| **Data collection and processing** | | | | |
| Microscope | Titan Krios G4 | Titan Krios G4 | Titan Krios G4 | Titan Krios G4 |
| Camera | Falcon 4i | Falcon 4i | Falcon 4i | Falcon 4i |
| Energy Filter | Selectris X (10 eV) | Selectris X (10 eV) | Selectris X (10 eV) | Selectris X (10 eV) |
| Magnification | 165'000 | 165'000 | 165'000 | 165'000 |
| Voltage (kV) | 300 | 300 | 300 | 300 |
| Electron exposure ($e^{-}/Å^2$) | 40.0 | 40.0 | 40.0 | 40.0 |
| Defocus range (μm) | -0.8 to -2.5 | -0.8 to -2.5 | -0.8 to -2.5 | -0.8 to -2.5 |
| Pixel size (Å) | 0.732 | 0.732 | 0.732 | 0.732 |
| Symmetry imposed | $C_3$ | $C_3$ | $C_3$ | $C_3$ |
| Initial particle images (no.) | 363’408 | 1’352’936 | 2’394’673 | 1’602’206 |
| Final particle images (no.) | 50’000 | 50’000 | 50’000 | 50’000 |
| Map resolution (Å) | (2.9) | (2.8) | (3.0) | (3.0) |
| FSC threshold | 0.143 | 0.143 | 0.143 | 0.143 |
| Map sharpening *B* factor ($Å^2$) | 73.1 | 60.2 | 68.1 | 75.6 |
| Orientation Diagnostics | | | | |
| cFAR | 0.02 | 0.02 | 0.03 | 0.01 |
| SCF* | 0.60 | 0.56 | 0.58 | 0.55 |

### 7 | References


1. Punjani, A., Rubinstein, J. L., Fleet, D. J. & Brubaker, M. A. cryoSPARC: algorithms for rapid unsupervised cryo-EM structure determination. *Nat. Methods* **14**, 290–296 (2017).
2. Krüger, C. R., Mowry, N. J., Bongiovanni, G., Drabbels, M. & Lorenz, U. J. Electron diffraction of deeply supercooled water in no man’s land. *Nat. Commun.* **14**, 2812 (2023).
3. Mowry, N. J., Krüger, C. R., Bongiovanni, G., Drabbels, M. & Lorenz, U. J. Flash melting amorphous ice. *J. Chem. Phys.* **160**, 184502 (2024).
